# HLiCA: An integrated cell atlas of the healthy human liver

**DOI:** 10.64898/2026.06.30.735539

**Authors:** Rachel D. Edgar, Jordan R. Portman, Hongru Hu, Delaram Pouyabahar, Raza R. Rahman, Daniel Stueckmann, Yongin Choi, Drew R. Neavin, Jawairia Atif, Zoe A. Clarke, Ran Gao, Shruti Khare, Ziyi Li, Liesbet Martens, Abhishek Murti, Diana Nakib, Niranjan Shirgaonkar, Stefan Thomann, Tinne Thoné, John R. Wilson-Kanamori, Katja Breitkopf-Heinlein, Elias Isaac Lattouf, Ruoxin Li, Renee Napoliello, Nuh N. Rahbari, Mehrshad Sadria, Oran Yakubovsky, Tallulah Andrews, Bruce J. Aronow, Alex G. Cuenca, Erica A.K. DePasquale, Stacey S. Huppert, Shalev Itzkovitz, Georg M. Lauer, Krupa R. Mysore, Joseph E. Powell, Robert E. Schwartz, Ankur Sharma, Sarah A. Taylor, Ludovic Vallier, Bruce Wang, Ramanuj Dasgupta, Dominic Grün, Martin Guilliams, Neil C. Henderson, Sonya A. MacParland, Charlotte L. Scott, Alan Mullen, Gerald Quon, Gary D. Bader

**Affiliations:** Ajmera Transplant Centre, Toronto General Research Institute, University Health Network, Toronto, Ontario, Canada; Donnelly Centre, University of Toronto, Toronto, Ontario, Canada; Centre for Inflammation Research, Institute for Regeneration and Repair, University of Edinburgh, Edinburgh, UK; Department of Molecular and Cellular Biology, Genome Center, University of California, Davis, CA, USA; Department of Molecular Genetics, University of Toronto, Toronto, Ontario, Canada; Eric and Wendy Schmidt Center, Broad Institute of MIT and Harvard, Cambridge, MA, USA; Division of Gastroenterology, University of Massachusetts Chan Medical School, USA; Broad Institute of MIT and Harvard, Cambridge, MA, USA; Department of Biomedical Engineering, University of California, Davis, CA, USA; Centre for Population and Disease Genomics, Institute for Molecular Bioscience, University of Queensland, Brisbane, Australia; Translational Genomics Program, Garvan Institute of Medical Research, Darlinghurst, NSW, Australia; Department of Immunology, University of Toronto, Toronto, Ontario, Canada; School of Cancer Sciences, University of Glasgow, Glasgow, UK; Cancer Research UK Scotland Institute, Glasgow, UK; National Key Laboratory of Immunity and Inflammation, Suzhou Institute of Systems Medicine, Chinese Academy of Medical Sciences & Peking Union Medical College, Suzhou, China; Laboratory of Myeloid Cell Biology in Tissue Damage and Inflammation, VIB-UGent Center for Inflammation Research, Ghent, Belgium; Causal Systems Immunology Laboratory, VIB-UGent Center for Inflammation Research, Ghent, Belgium; Department of Biomedical Molecular Biology, Faculty of Sciences, Ghent University, Ghent, Belgium; Department of Medicine, Division of Gastroenterology, University of California San Francisco, CA, USA; Würzburg Institute of Systems Immunology, Julius-Maximilians-Universität Würzburg, Germany; Department of Surgery, Medical Faculty Mannheim, Heidelberg University, Mannheim, Germany; Gastrointestinal Division, Massachusetts General Hospital and Harvard Medical School, Boston, MA, USA; Department of Surgery, University Hospital Ulm, Ulm, Germany; Department of Applied Mathematics, University of Waterloo, Waterloo, Ontario, Canada; Department of Molecular Cell Biology, Weizmann Institute of Science, Rehovot, Israel; Department of Biochemistry, Schulich School of Medicine and Dentistry, and Department of Computer Science, University of Western Ontario, Canada; Divisions of Biomedical Informatics and Developmental Biology, Cincinnati Children’s Hospital Research Foundation, Cincinnati, OH, USA; Department of Computer Science, University of Cincinnati, Cincinnati, OH, USA; Department of Surgery, Boston Children’s Hospital, Boston, MA, USA; Division of Biomedical Informatics, Cincinnati Children’s Hospital Medical Center, Cincinnati, OH, USA; Division of Gastroenterology, Hepatology and Nutrition, Cincinnati Children’s Hospital Medical Center, Cincinnati, OH, USA; Department of Pediatrics, University of Cincinnati, Cincinnati, OH, USA; Division of Pediatric Gastroenterology, Hepatology and Nutrition, Texas Children’s Hospital, Baylor College of Medicine, Houston, TX, USA; William Shearer Center for Human Immunobiology, Texas Children’s Hospital, Houston, TX, USA; UNSW Cellular Genomics Futures Institute, University of New South Wales, Sydney, Australia; Division of Gastroenterology and Hepatology, Weill Cornell Medicine, New York, NY, USA; School of Clinical Medicine, Faculty of Medicine and Health, University of New South Wales, Sydney, Australia; KK Research Centre, KK Women’s and Children’s Hospital, Singapore; Department of Pediatrics, University of Colorado School of Medicine, Aurora, CO, USA; Berlin Institute of Health Centre for Regenerative Therapies (BIH@Charité), Berlin, Germany; Max Planck Institute for Molecular Genetics, Berlin, Germany; MRC Human Genetics Unit, Institute of Genetics and Cancer, University of Edinburgh, Edinburgh, UK; Department of Laboratory Medicine and Pathobiology, University of Toronto, Toronto, Ontario, Canada; Department of Computer Science, University of Toronto, Toronto, Ontario, Canada; Princess Margaret Cancer Centre, University Health Network, Toronto, Ontario, Canada; Lunenfeld-Tanenbaum Research Institute, Sinai Health System, Toronto, Ontario, Canada; CIFAR Multiscale Human Program, CIFAR, Toronto, Ontario, Canada

## Abstract

The human liver is composed of a heterogeneous mix of cell types. How these distinct populations contribute individually and collectively to liver function remains poorly understood. Although single-cell technologies have advanced our understanding of liver biology, individual studies have often been limited by small donor cohorts and inconsistent cell type annotations. Integrating multiple datasets can overcome these challenges and better capture biological variability.

We present the Human Liver Cell Atlas (HLiCA), an integrated reference of non-disease liver cells assembled from eight datasets across six research centers, encompassing more than 525,000 cells from 110 donors. Developed in collaboration with the Human Cell Atlas Liver Bionetwork, the HLiCA incorporates expert-curated cell annotations refined through community feedback and dedicated cell type annotation meetings. The HLiCA classifies cells into six lineages and expands the cell type resolution to include 47 distinct cell types. Starting from raw sequencing reads, we realigned all data and performed rigorous benchmarking to ensure robust integration across technical and biological variables. Genetic ancestry was inferred for all samples to evaluate the range of ancestral backgrounds represented in the atlas.

The expanded cell type annotation enabled identification of previously unrecognized liver cell types, including *NRXN1^+^* stromal cells. Their presence was validated using spatial transcriptomics, which localized *NRXN1^+^* stromal cells to periportal regions. With the number of donors included in the HLiCA we were able to examine cell type specific associations with demographic covariates. In hepatocytes, drug metabolism genes showed differential expression between sexes, and in cholangiocytes, mucus-production genes varied with age.

As the largest and most genetically diverse human liver cell atlas to date, the HLiCA provides a comprehensive, well-annotated reference for the field, annotated by expert consensus. This resource will enable deeper interrogation of liver cellular diversity, architecture, and function in the healthy human liver and serve as a reference to understand changes that occur with disease.

## INTRODUCTION

The human liver is a remarkable organ with a unique capacity for regeneration and a central role in maintaining health. The liver performs a wide range of physiological functions including vital roles in metabolism, blood detoxification, protein synthesis, and immune regulation. These crucial functions are accomplished by a complex interplay between cell types. Much of our understanding of the specialized cellular composition of the healthy liver comes from single-cell and single-nucleus RNA-sequencing (sc/snRNA-seq) maps^1–9^. Often, these maps of the healthy human liver are generated as a comparator for studying liver disease, such as primary sclerosing cholangitis^2^, metabolic dysfunction-associated steatotic liver disease^10^, hepatitis B^5^, fibrosis^4^ or cirrhosis^11^, while other studies have focused on understanding the liver in health and have generated maps of the healthy liver without a specific disease comparison^2,3,6^.

Whether characterizing the healthy or diseased liver, sc/snRNA-seq studies are typically tailored to the underlying research focus which can limit their use toward other research questions. For example, experimental protocols may intentionally enrich specific cell types^6,11^ or process tissue in ways optimized to preserve some cell types at the expense of others^1^. The broad applicability of sc/snRNA-seq maps is also limited by available cohort sizes. Existing maps are limited to donor cohorts available at individual research centres and are currently not large enough to explore how demographic factors such as age, sex, and genetic ancestry influence liver cell populations. Similarly, studies from individual centers may lack the cell numbers required to characterize and confidently classify rare cell populations increasing the probability that the full cellular diversity of the healthy liver has not yet been described. The limited cohort sizes and cell numbers combined with a variety of experimental protocols used means no one existing liver map is tailored to serve as a universal reference.

Integrating multiple maps into a combined atlas can overcome these limitations and is an approach already used for several tissues as part of the Human Cell Atlas (HCA)^12–14^. These cross-cohort integrated atlases have led to the characterization of newly identified cell types and markers, as well as associations between donor demographic information and gene expression^13–16^. With these existing tissue atlases serving as a model, but adapted for the unique challenges of the human liver, a roadmap for building a healthy human liver atlas was established^17^. Following this roadmap we have assembled the Human Liver Cell Atlas (HLiCA), an integrated atlas of over 525,000 non-diseased human liver cells from 110 donors. The assembly of the HLiCA involved the collaboration across the HCA Liver Bionetwork, and has led to the characterization of newly identified cell types and associations of gene expression with age and sex. Liver tissue has defined spatial structure and hepatocyte zonation is one of the strongest gene expression patterns in the liver^18,19^. We therefore extended the atlas to spatial transcriptomic data and explored the newly identified cell types and demographic associations spatially.

This work represents an initial version of a human liver cell atlas, designed to serve both as a high-quality reference resource for the field but also as a foundation to be built upon as additional data become available. As the largest and most genetically diverse human liver cell atlas generated to date, the HLiCA enables an in-depth investigation of liver cellular diversity, spatial organization, and functional states in the healthy human liver.

## METHODS

### Data collection from participating studies

Data from seven landmark studies were included in HLiCA^1–6,9^ and only the samples derived from healthy donors in those studies were included (Table S1). We focused our data collection on studies that used 10x Genomics single-cell and single-nucleus RNA-seq (sc/snRNA-seq) to reduce technology-related variability. To minimize technical artifacts arising from differences in reference genome or alignment software versions, we aimed to realign all samples and therefore only included studies with publicly available raw sequencing data (FASTQs). For the previously unpublished data presented here, all FASTQs have been made available (see Data Availability Statement). Details on data generation of the previously unpublished samples is provided in the supplementary materials (Table S2-3). Ethics approval details for published data are provided in original publications^1–6,9^ and for unpublished data details are in the Supplementary Methods.

### Metadata curation and harmonization

Metadata for all samples (donors/sequencing libraries) were curated and harmonized to ensure consistency across datasets (Supplementary Data 1-2). The key technical covariates included ‘STUDY’, ‘suspension_type’, ‘assay’, ‘donor_uuid’, and ‘library_alias’. This harmonized metadata facilitated downstream integration and analysis processes.

### Sequence alignment and count matrix generation

The FASTQ files from collected studies were aligned to the reference genome using Cell Ranger v7.0.0. The reference human genome was downloaded from the GENCODE Release 42 (GRCh38.p13) repository (gencodegenes.org/human/release_42.html) and was used for sequence alignment. Each dataset was processed through Cell Ranger with default parameters to produce the count matrices.

### Quality control

Quality control (QC) was performed on each count matrix using the pipeline described previously^2^. Briefly, to distinguish true cells from empty droplets, droplet filtering was performed using an EmptyDrops with a false discovery rate (FDR) threshold of 0.01^20^. Droplets with total counts below 10 unique molecular identifiers (UMIs) were excluded. Cells were required to express at least 100 genes and to have mitochondrial gene expression below 50%. Genes were retained if detected in at least 10 cells. Doublets were excluded during the clustering and cell annotation process described below.

### Further data selection

Additional steps were taken to refine the dataset. Blood samples included in one study were removed to focus on liver cells^5^. Duplicated sequencing runs published in multiple studies were identified and excluded to prevent redundancy in the HLiCA^1–3^. The final merged dataset included 33,670 genes and 644,680 cells, with 156 sequencing runs/libraries from 110 donors.

### Data normalization

Following initial QC, the standard Seurat preprocessing pipeline was applied for normalization, variable feature identification, and scaling^21–24^. The NormalizeData function was used to normalize the gene expression measurements for each cell by the total expression counts, multiplying by a scale factor (10,000), and log-transforming the result. Variable features were determined using the ‘vst’ method via FindVariableFeatures function, with 4,000 highly variable features selected.

### Integration evaluation

In order to select the most appropriate integration method for HLiCA we evaluated four methods: Harmony^25^, scVI^26^, Seurat rPCA^22^ and Weighted Nearest Neighbours (WNN) ^22^. Integration performance was assessed for three metadata variables: sample ID, suspension type and a reference based cell type prediction from scmap^27^ which was performed on the whole atlas prior to any integration to avoid bias to cell types derived from an integrated embedding. Integration was evaluated with two metrics: Adjusted Rank Index (ARI) (R: mclust v6.1.1)^28^ and Local Inverse Simpson Index (LISI) (R: lisi v1.0)^25^.

### Data integration with Harmony

Data was integrated using Harmony (v0.1.1)^25^ to correct for batch effects and technical variation across datasets. Specifically, Harmony was run with the following parameters: group.by.vars set to the covariates list (‘STUDY’, ‘suspension_type’, ‘assay’, ‘donor_uuid’, and ‘library_alias’), assay.use set to ‘RNA’, and reduction set to ‘pca’ with dims set to the first 40 principal components. Subsequently, the integrated latent embedding generated by Harmony was used for downstream analysis, including cell clustering and visualization to aid cell annotation.

### Cell annotation

To define liver cell types, we first clustered (Louvain clustering, resolution 0.5, 40 dimensions) cells in the entire Harmony integrated map into 6 lineages (Alpha Annotation: Hepatocyte, Lymphocyte, Endothelia, Myeloid, Mesenchyme, and Cholangiocyte) based on known marker gene expression (Supplementary Data 3). Then, within each lineage, we integrated with Harmony and the same covariates used in the entire atlas integration. This integration and clustering within the lineages formed the initial draft annotation of cell types which was based on lineage specific cell type markers (Supplementary Data 4), as well as CIPR used on the 6 lineages^29^. These initial draft annotations were then evaluated in a series of cell type annotation meetings to discuss cell identities and marker expression. At each phase of the annotation (lineage annotation, initial draft annotation and final HLiCA annotation) we pruned suspected doublets based on cluster level expression of markers from multiple lineages (Supplementary Data 3). After doublet removal, another round of within lineage Harmony integration and clustering was performed, with the final HLiCA cell type labels based on these clusters and the feedback from the cell type annotation meetings.

To characterize identities and potential functions of cell types, we performed differential gene expression analysis across all cell types within each of the six major lineages. For each lineage, every cell type was compared against all other cell types using the Seurat function *FindAllMarkers*^21–24^ with default settings. The resulting fold-change values were then used to assess pathway enrichment for each cell type via fGSEA^30^, using pathways from the Enrichment Map Gene Sets (Human Gene Ontology, including inferred electronic annotation, October 26, 2022 release)^31^.

To evaluate how the HLiCA cell type annotations compared with those provided in the original publications for these data, we generated confusion matrices between the original annotations and our HLiCA annotations for the same cells. Specifically, for each cell type annotated in the HLiCA, we calculated the proportion of cells assigned to each of the original authors’ annotation cell types. This analysis was restricted to studies with published cell annotations and limited to cells passing quality control in both our analysis and the original analysis.

### Ancestry admixture estimation

Genetic admixture was estimated using the genetic information captured from the sc/snRNA-seq reads. First, variants were called from the sc/snRNA-seq reads with an altered form of Monopogen^32^ that only considers known common variant locations to reduce compute time and resources. This includes variant calling using samtools and bcftools followed^33^ by genotype imputation of the called variants with Beagle^34^. Donor admixture was estimated using the genotype likelihoods with fastNGSadmix^35^. A reference of allele frequencies was generated from the Human Genome Diversity Project (HGDP)^36^ after removing donors with more than 5% admixture from other continental regions. Many individuals in the HLiCA show genetic similarity to multiple HGDP populations. To summarize genetic ancestry at the cohort level, we defined primary estimated ancestry as the HGDP population with the largest estimated proportion in each individual (range largest proportions: 0.4–0.9999).

### Differential expression analysis

To investigate associations between gene expression, age, and sex in the HLiCA, we applied a linear mixed model using FLASH-MM^37^. This framework enabled us to account for library (i.e., sample) as a random effect, which should reduce the p-value inflation often observed in sc/snRNA-seq differential expression analysis^38,39^. For both age and sex models, library was modeled as a random effect, with study and suspension type included as fixed effects in the sex model. Genes were considered significantly differentially expressed if they had a FDR < 0.05 and an absolute coefficient for the sex or age term > 0.5. Consistent with the pathway analysis performed for newly identified cell types, we again used fGSEA^30^ with the same gene sets, but genes were ranked by their coefficients for either sex or age. We excluded two samples from the sex and age analyses because their sex chromosome gene expression did not match the reported sex. An additional sample, which did not conform to either male or female sex chromosome expression patterns, was removed from the sex analysis but kept in the age analysis (Supplementary Methods).

### Validation of cell types in spatial transcriptomics

To validate the existence of newly identified cell types we used available spatial transcriptomics data from healthy human adult livers. Specifically three samples from three donors assayed using a 317 gene MERFISH panel^4^, 4 samples from 3 donors assayed using a 477 gene Xenium panel^40^, and one sample assayed using Visium HD^41^. For these three datasets the published cell type labels of segmented cells or bins was used, with further subclustering of relevant cell types using marker genes of the newly identified cell types discovered in the HLiCA. For the quantification of zonation of *NRXN1*^+^ stromal cells in Xenium data, previously published distances from periportal and pericentral regions were used^40^. The zonation metric shown is the difference between the distance to the nearest periportal region minus the distance to the nearest pericentral, as cells could be near multiple vascular features. In the Visium HD spatially adjacent cholangiocyte bins were grouped using a graph-based connected components algorithm to define bile ducts, as previously described^41^. Bile ducts were classified by diameter according to established duct size categories^42^. Septal ducts had a mean diameter of 147 µm (range 101–209 µm), and interlobular ducts had a mean diameter of 54 µm (range 23–93 µm). Bile ductules were defined as <20 µm in diameter and corresponded to ducts measuring 16 µm in our dataset (two 8 µm spatial bins).

## RESULTS

### Data integration establishes the healthy human liver atlas

To create the HLiCA, we assembled single-cell RNA sequencing (scRNA-seq) and single-nucleus RNA-seq (snRNA-seq) data from eight datasets (seven published and one unpublished), across six research centres (Fig. 1AB, Table S1). These datasets include samples from 110 individuals with diversity in age and genetic ancestry (Fig. 1CE). Cells were obtained from 156 tissue samples using a variety of sampling methods and experimental protocols (Fig. 1D, Supplementary Data 1-2, Supplementary Methods). After quality control, the atlas contained 524,699 cells integrated across studies. Our integration approach was chosen after benchmarking several alternates (Fig. S1; Supplementary Results), and we observed reasonable integration across assay, research center and suspension type (scRNA-seq vs snRNA-seq) (Fig. S2).

**Figure 1:**
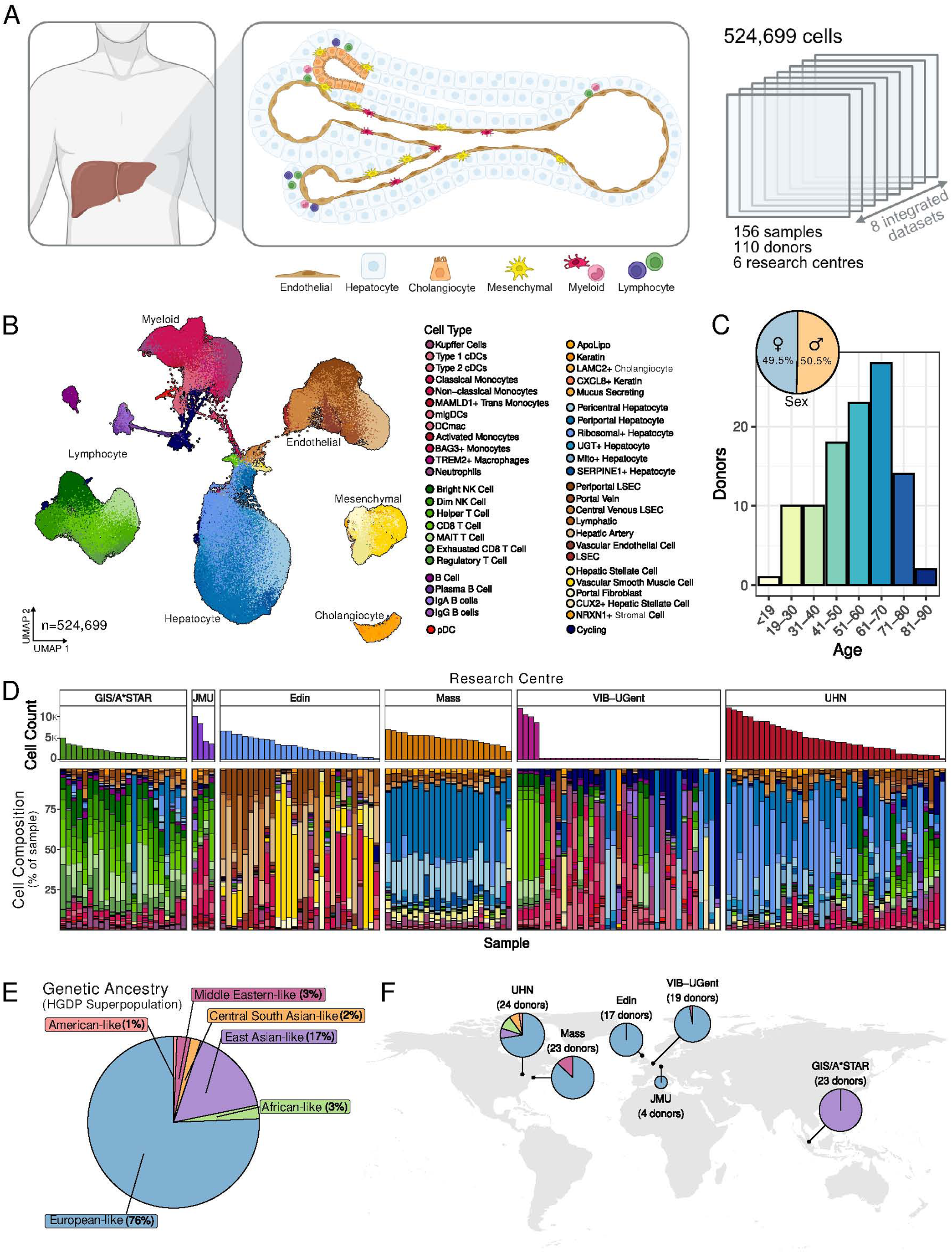
Overview of the HLiCA. A) A schematic overview of the cell types we aimed to capture in the atlas and the datasets included in the integration. B) UMAP of the integrated HLiCA coloured by the final cell annotation. C) Distribution of ages and sex in the HLiCA. D) Cell composition of each sample in HLiCA, split by Research Centre. E-F) Proportion of genomes similar to each HGDP superpopulation E) across the entire HLICA F) split by the Research Centre.

Cell type annotation was an iterative process. Initially we grouped cells into six lineages based on clustering and canonical makers (Fig. S3-5; Table S6). Through this process we annotated 47 cell types in HLiCA (Fig. 1BD). Cell type-specific differentially expressed genes (FDR < 0.005 and |log_2_ fold change| > 1) and corresponding gene set enrichment analyses are provided in Supplementary Data 5-6. In line with ongoing efforts to standardize liver cell nomenclature, we incorporated HLiCA cell types into the Cell Ontology (CL; Supplementary Data 7). Despite differences in sampling procedures between research centres (Fig. 1D, Table S1) all but two cell types were well represented from multiple data centers (Fig. S6), and we observed an enrichment of myeloid and lymphocyte cells in scRNA-seq compared to snRNA-seq, as previously reported (Fig. S7)^1^. Plasmacytoid dendritic cells (pDCs) are included in the entire atlas but not in any lineage maps as they are transcriptomically distinct from both the myeloid lineage and lymphocyte lineage (Fig. S5).

Although we did not use previously published cell labels during our annotation process, we compared our annotations to these labels to ensure our annotation was aligned with previous work. Overall, the HLiCA annotations had an expected overlap with prior labels but with greater granularity in all lineages (Fig. S8). For example, Regulatory T cells were identified across research centres in the HLiCA dataset (Fig. 2A), even though this population was not annotated in most of the original studies (Fig. 2B). Where high-resolution lymphocyte annotations were previously available, such as in the NK and T cell focused GIS/A*STAR datasets, we observed strong correspondence between our Regulatory T cell label and the previously defined “Treg” populations (Fig. 2C). The HLiCA can be used as a reference to identify rare cell populations, like regulatory T cells, in smaller maps such as the healthy human pediatric liver^40^ (Supplementary Results; Fig. S9). By integrating multiple datasets into a combined atlas, we provide improved granularity across lineages, enabling characterization of populations such as Regulatory T cells and increasing our confidence in their identification in individual datasets with smaller cohort sizes.

**Figure 2:**
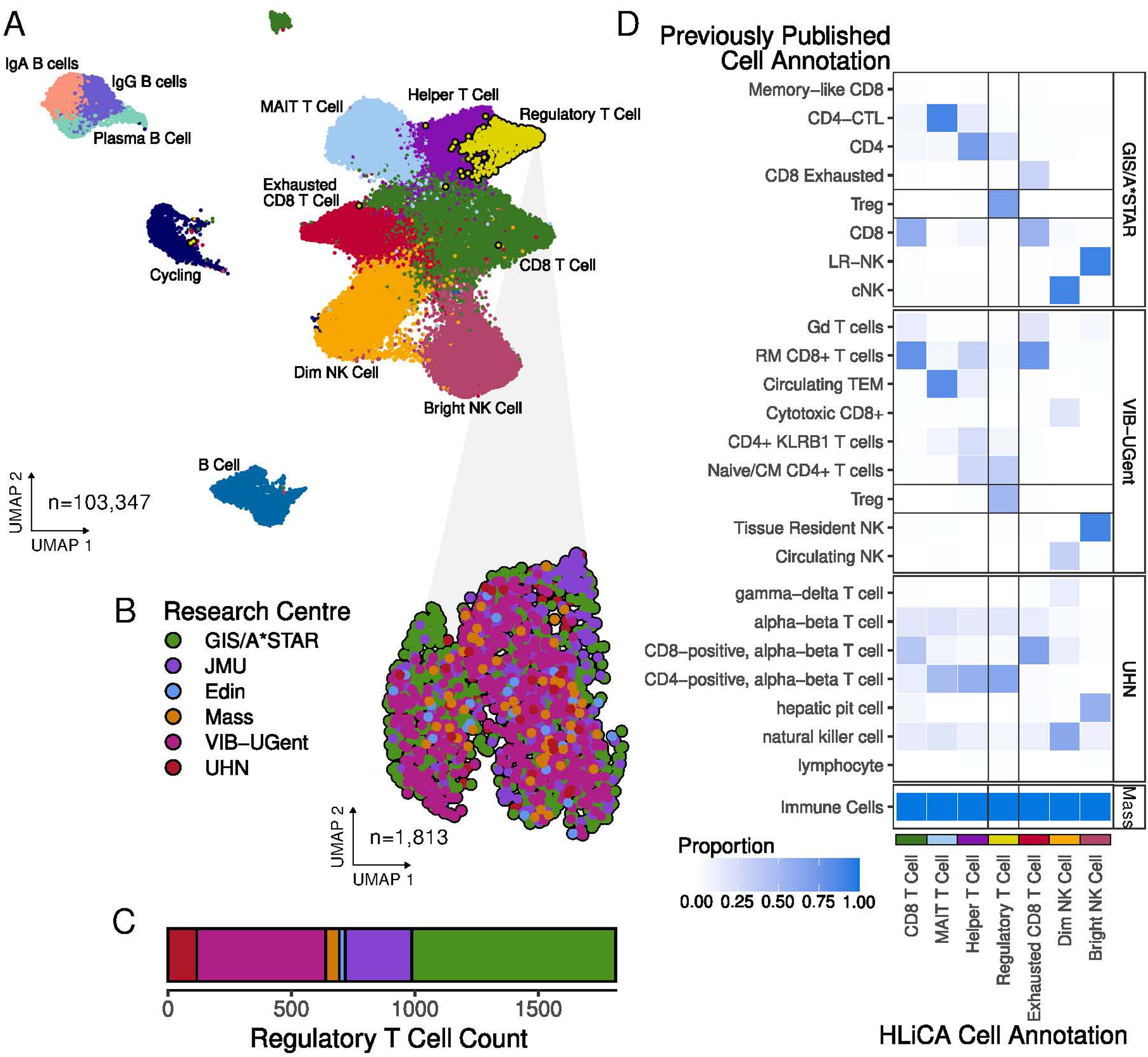
Refined cell-type annotation identifies regulatory T cells across centres. A) UMAP of 12 lymphocyte subtypes with regulatory T cells highlighted. B) UMAP of only regulatory T cells coloured by Research Centre. C) Cell counts of regulatory T cells from each research centre. D) Confusion matrix between published annotation and the HLiCA Gamma annotation of lymphocytes.

### Rare and newly identified cell types

#### Liver myeloid cell landscape

We identified a subset of monocytes characterized by high expression of *MAMLD1*, which resemble a previously described population of cells transitioning from monocyte-to-macrophage in the mouse liver (*Clec4e^+^* macrophages)^43^. Notably, our *MAMLD1*^+^ trans monocytes uniquely express high levels of *CXCL2*, a marker associated with this transitional state (Supplementary Data 5). This population is similar to the monocyte subset described in models of metabolic dysfunction-associated steatohepatitis (MASH), representing a recruited monocyte state that either progresses toward macrophage differentiation or undergoes cell death^43^. The presence of this population in healthy liver may reflect a low-level, recruitment and differentiation of monocytes into macrophages that contribute to steady-state macrophage turnover or surveillance. We also identified *TREM2*^+^ macrophages that, together with the *MAMLD1*^+^ trans monocytes, share gene expression signatures similar to the previously described lipid-associated macrophage (LAM)-like populations^6,44^ (Supplementary Data 5). Furthermore, we observed a population of migratory dendritic cells (migDCs) that express *CCR7*, *LAMP3*, and *BIRC3*, which are transcriptionally similar to cells termed DC3 in some studies^45,46^ (Supplementary Data 5). These findings support the presence of diverse, functionally specialized myeloid subsets in the healthy liver, some of which mirror populations previously associated with inflammatory or metabolic disease states.

#### Liver mesenchyme cell landscape

We also identified a rare cell population (n=347) that clustered with mesenchymal lineage cells, which we have labeled *NRXN1*^+^ stromal cells. These *NRXN1*^+^ stromal cells were not previously resolved in individual liver maps (Fig. 3A-E). These cells exhibit gene expression patterns and pathways related to the “*Ensheathment of Neuro*ns” (Table SX). Using spatial transcriptomics we observed a similar population of *NRXN1*^+^ stromal cells, enriched in periportal regions (Fig. 3F, S10). As with regulatory T cells, the HLiCA annotation has enhanced the ability to resolve rare cell populations. Once defined, *NRXN1*^+^ stromal cells could be observed in the healthy pediatric liver, but only when the pediatric map was combined with the HLiCA (Supplementary Results; Fig. S11).

**Figure 3:**
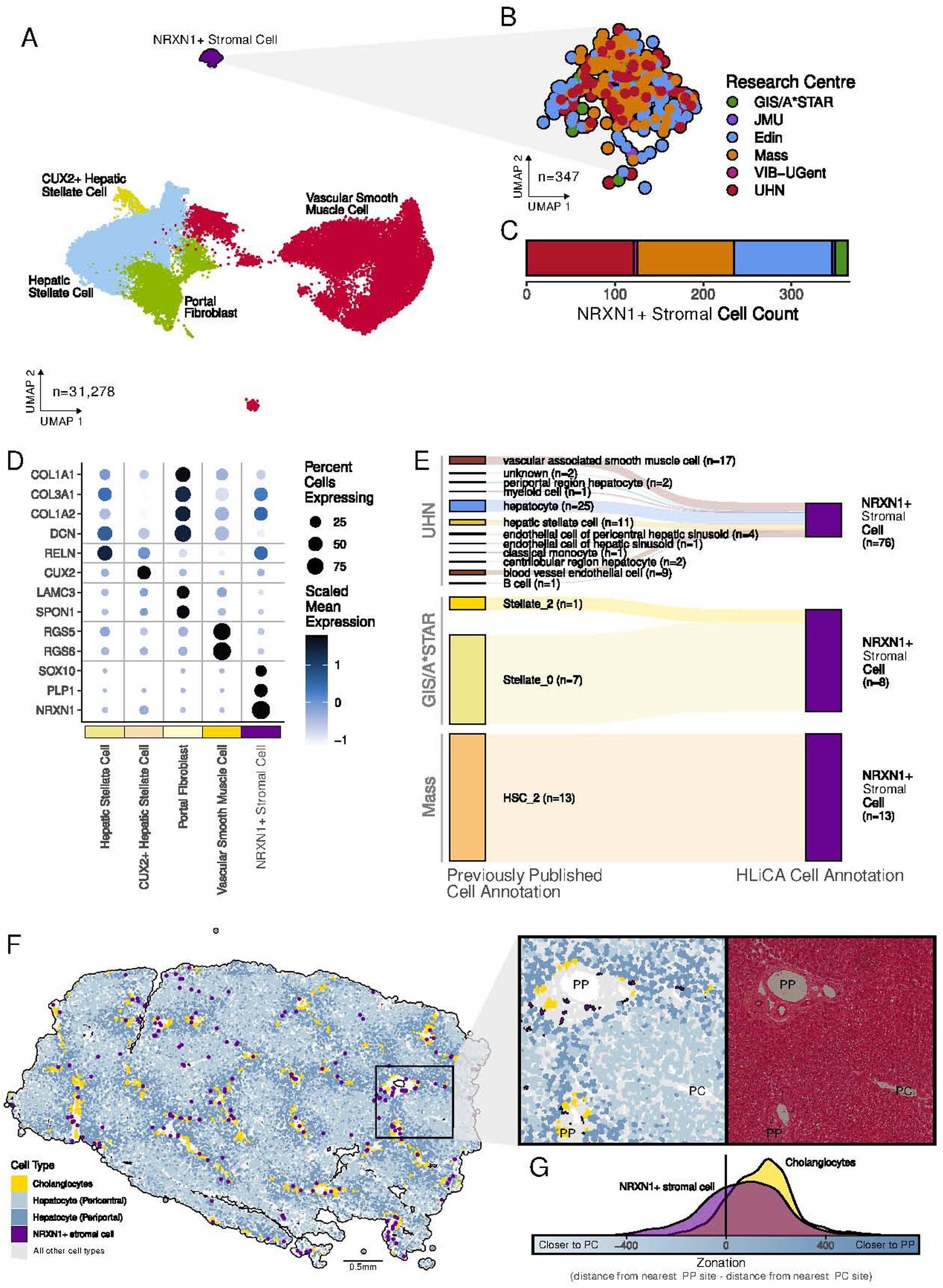
NRXN1+ stromal cells were captured by all research centres and are more abundant in periportal regions. A) UMAP of mesenchymal lineage with NRXN1+ stromal cells highlighted. B) UMAP of only NRXN1+ stromal cells coloured by Research Centre. C) Cell counts of NRXN1+ stromal cells from each research centre. D) Expression of mesenchymal marker genes. E) Annotation of the NRXN1+ stromal cells in the previous publications with these data. F) Location of a population of NRXN1+ stromal cells in Xenium spatial transcriptomics. Only selected cell type annotations are shown to highlight zonation patterns. Cell segmentation is shown along side H&E staining in the zoomed in panel. G) Zonation of NRXN1+ stromal cell and cholangiocytes across Xenium samples (n=4).

#### Liver cholangiocyte cell landscape

Mucus-secreting cholangiocytes characterized by high expression of secretory and antimicrobial genes were also defined through the HLiCA (Fig. 4A-C). This population of cholangiocytes exhibit upregulation of *MUC5B, MUC13, MUC1, MUC3A, and MUC12*, supporting their classification as mucus-producing cells (Supplementary Data 5). Trefoil factors (TFF1, TFF2, and TFF3), which play roles in mucosal repair^47,48^, and related mucus production proteins such as CEACAM6, SCGB2A1, and SCGB3A1 were also enriched. These cells showed pathway enrichment for epithelial cell differentiation and mucus secretory functions, including “*Keratinization*”, “*O-Linked Glycosylation of Mucins*”, as well as antimicrobial functions such as “*Neutrophil Degranulation*”, and “*Antimicrobial Humoral Response*” (Supplementary Data 6). In spatial transcriptomics, we were able to identify 978 mucus secreting cholangiocytes located, as expected, in periportal regions (Fig. 4D). When bile ducts were binned by diameter, we observed mucus secreting cholangiocytes were found in bile ducts of all diameter sizes but more frequently in larger bile ducts (Fig. 4EF, chi-squared p<0.05). Although similar mucus-secreting cholangiocytes have been previously observed^2,49^, by integrating data across studies, we were able to define a larger set (n = 471) of these relatively rare cells.

**Figure 4:**
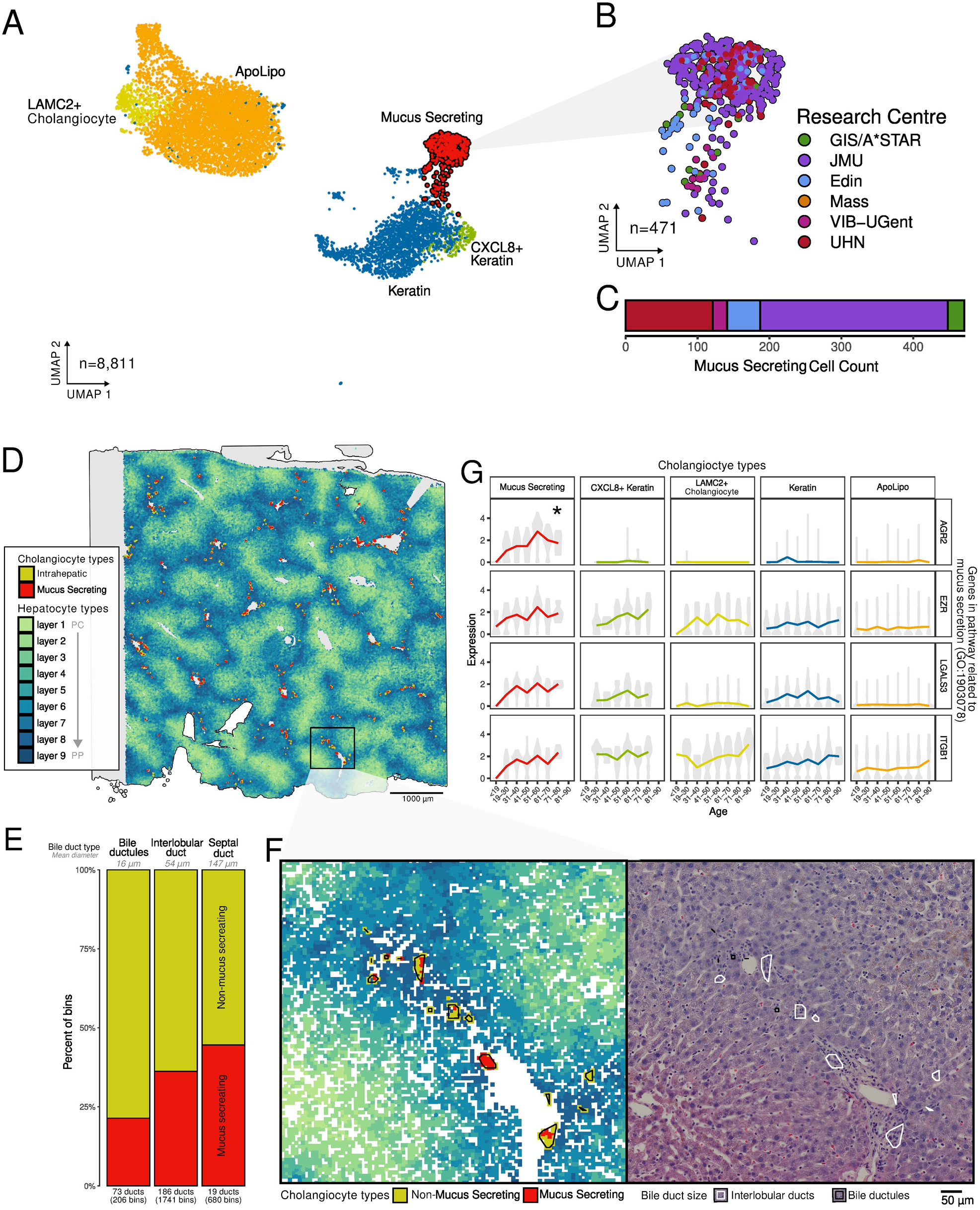
Gene expression in mucus secreting cholangiocytes changes with age. A) UMAP of cholangiocyte lineage with mucus secreting cholangiocytes highlighted. Points are coloured by cell type. B) UMAP of only mucus secreting cholangiocytes coloured by Research Centre C) Cell counts of mucus secreting cholangiocytes from each research centre. D) Mucus secreting cholangiocytes are more frequently near periportal regions than pericentral regions. E) Cholangiocyte composition of bile ducts split by bile duct size. F) Representative periportal region enlarged with bile ducts outlined in cell annotation view. Paired H&E staining shows bile ducts outlined an coloured by size category. G) Genes driving the enrichment of mucus secretion pathways show increasing expression with age mainly in Mucus secreting cholangiocytes.

#### Liver hepatocyte cell landscape

The hepatocyte lineage was challenging to annotate, in part due to the presence of several populations that may originate from technical artifacts but could represent true biological variation (e.g., Mito^+^ and Ribosomal^+^ hepatocytes; Fig. 5A, S5). With that caveat in mind, we observed the expected pericentral and periportal hepatocyte populations (Fig. 5A, S5). However, we were unable to resolve a distinct mid-zonal population, likely due to the limited availability of well-established transcriptional markers for these cells. Despite this limitation, we defined previously uncharacterized hepatocyte subsets with differentially expressed genes and pathways suggesting possible functional roles in the healthy human liver. We identified a population of *SERPINE1^+^* hepatocytes, which was characterized by high *SERPINE1* expression and enrichment of transcripts encoding genes in the TGF-β signaling pathway (Fig. 5A, S5; Supplementary Data 5-6). Notably, TGF-β signaling has been reported to play roles in both cellular senescence^50^ and lipid metabolism^51^. This subset also showed increased expression of hepatocyte injury-associated genes including *KLF6*, *SOX4*, *DUSP8*, *BIRC3*, and *TGFB* and was enriched for the “*Apoptosis Signaling Pathway,*” suggesting a potential role in liver injury or cellular stress responses. *UGT*^+^ hepatocytes are another previously uncharacterized hepatocyte population (Fig. 5AB). Although this subset showed relatively few upregulated genes, it was enriched for multiple drug metabolism pathways, including “*Aspirin Absorption, Distribution, Metabolism and Excretion (ADME)*,” “*Ibuprofen Action Pathway*,” and “*Metapathway Biotransformation Phase I and II*”, suggesting this population may represent a specialized hepatocyte subset involved in drug metabolism (Supplementary Data 5-6). Given the established role of pericentral hepatocytes in drug metabolism, we confirmed that the unique expression of UGT genes and the enrichment of drug-metabolizing pathways in UGT*^+^* hepatocytes remained greater when compared specifically to pericentral hepatocytes (Fig. S12). Using VisiumHD and MERFISH spatial transcriptomics datasets, we found that markers of *UGT^+^* hepatocytes were more highly expressed pericentrally (Fig. 5C-D).

**Figure 5:**
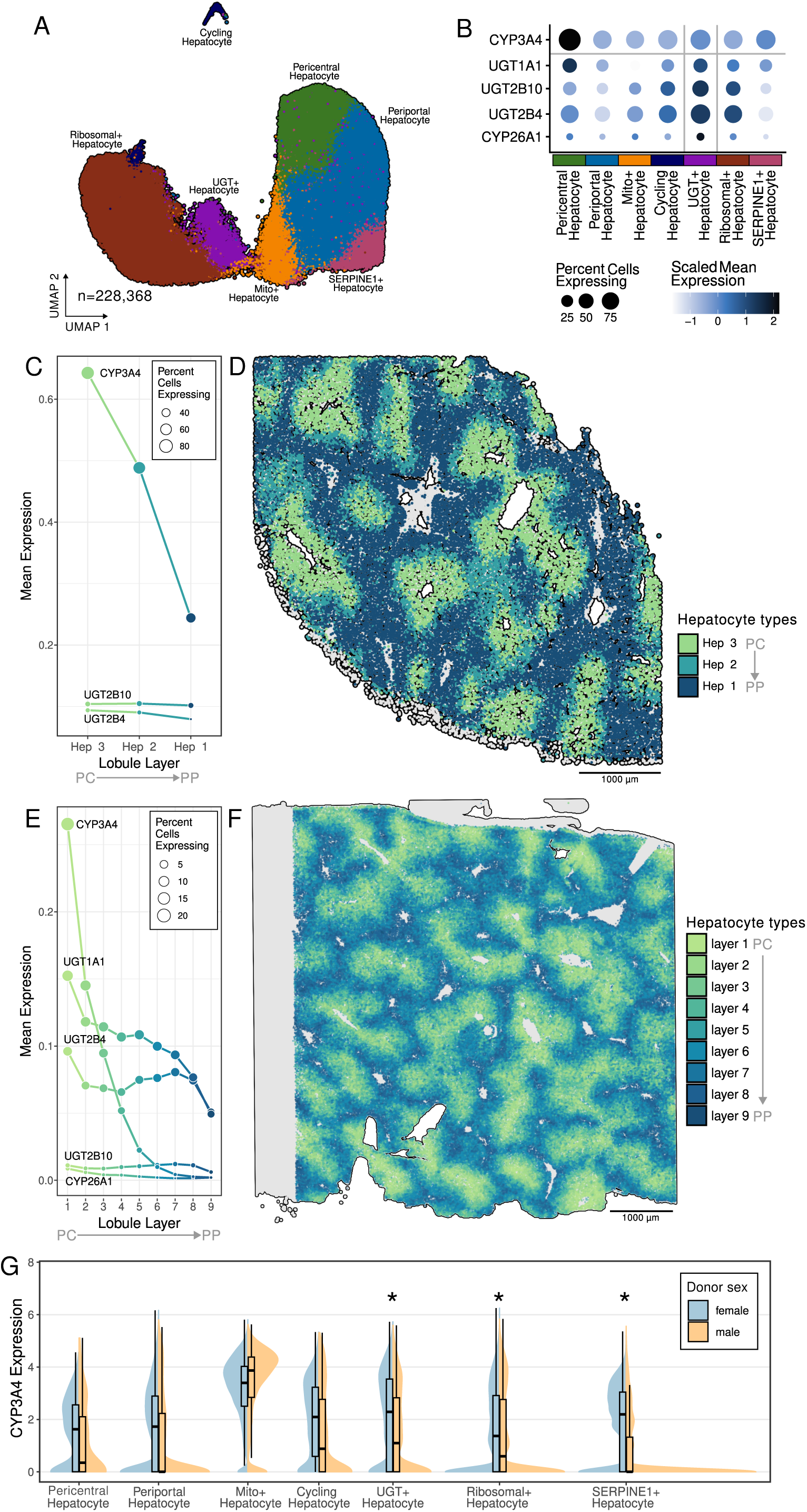
Female UGT+ hepatocytes show higher expression of CYP3A4. A) UMAP of hepatocyte lineage. B) Expression of UGT+ hepatocyte marker genes as well as CYP3A4. C) UGT genes and CYP3A4 show zonated expression with more expression in pericentral regions. D) Zonation of hepatocytes is captured with MERFISH spatial transcriptomics (n=3). E) UGT genes as a well as CYP3A4 are more highly expressed in pericentral regions (n=1). F) Zonation of hepatocytes is captured with VisiumHD spatial transcriptomics. G) Differential expression of CYP3A4 between sexes is significant in three hepatocyte cell types.

### Donor factors impact transcriptomic profiles

We observed differential expression with age in some populations (Fig. S13,14; Supplementary Data 8). Of particular interest was the increasing expression of *AGR2* with age in the mucus secreting cholangiocytes, together with enrichment of mucus secretion pathways among age associated genes (Fig. 4G; Supplementary Data 9). *AGR2* is known to be essential for mucus production in the intestine^52,53^ and mucus production has been shown to change with age^54^. The change in gene expression with age could suggest the regulation of mucus production changes with age in this relatively rare cholangiocyte population. We also observed differential gene expression between sexes in the newly identified *UGT*^+^ hepatocyte population (Supplementary Data 10-11). Notably, *CYP3A4* was more highly expressed in females *UGT*^+^ hepatocytes, as well as in *SERPINE1*^+^ and Ribosomal^+^ hepatocytes (Fig. 5G). *CYP3A4* is more highly expressed in females and is associated with faster clearance of drugs such as aspirin and paracetamol^55,56^. As *CYP3A4* and markers of *UGT^+^* hepatocytes are more highly expressed pericentrally, potentially the observed sex-differences in drug metabolism pathways may have the greatest effect at pericentral regions.

## DISCUSSION

### The HLiCA represents a comprehensive reference map of the healthy human liver, integrating transcriptomic profiles from eight datasets across six research centers and encompassing broad diversity in genetic ancestry, age, and sex

Through collaborative efforts within the HCA Liver Bionetwork, this resource captures extensive cellular diversity expanding the catalog of liver cell types. By incorporating spatial transcriptomic data, the HLiCA further contextualizes cellular identity within the structure of the liver. The representation of donor demographics positions the HLiCA as the most extensive and genetically diverse atlas of the healthy human liver to date. We anticipate the HLiCA will serve as a reference to advance understanding of liver biology in future studies. The HLiCA is accessible through multiple interactive platforms, including CELLxGENE, the Cell Annotation Platform (CAP) and the HCA data portal, enabling broad exploration and facilitating its use as a universal reference.

### The inclusion and reannotation of data across multiple centres revealed cell populations which were not characterized in individual maps

One rare and newly characterized cell type is the *NRXN1^+^* stromal cell populations. The genes that uniquely mark this cell type are related to the ensheathment of neurons, which is the function of Schwann cells as described in other tissues^57–61^. The physical location of *NRXN1^+^* stromal cells around periportal vasculature is consistent with the expected position of nerve fibres and their associated Schwann cells in the liver^62–65^. The evidence for the existence of Schwann cells in the liver is limited, but they are reported to be marked by the S-100 protein^62^ and their existence is supported by several case reports of hepatic schwannomas^66,67^. This is the first characterization of *NRXN1^+^* stromal cells using sc/snRNA-seq and the markers and annotation processes described here will allow for identification of *NRXN1^+^* stromal cells in future studies and the study of this unique population in relation to liver disease.

### With the number of donors included in the HLiCA, we were also able to identify gene expression differences associated with age and sex

The higher expression of *CYP3A4* in females is well known from studies of bulk liver samples^55^. Here we were able to narrow down the differential expression of *CYP3A4* to types of hepatocytes not previously described, such as the *UGT^+^* hepatocytes which have upregulated pathways related to drug metabolism. Importantly, we saw markers for the *UGT^+^* hepatocytes are more highly expressed in central veins, suggesting these hepatocytes are located toward central veins, which are key regions for drug metabolism^68^. The *UGT^+^*hepatocytes may be responsible, in part, for the differences in drug metabolism between men and women^56,69^. Given the limited clinical information available for many samples, the observed UGT^+^ hepatocyte population could reflect inter-individual variation in bilirubin metabolism, consistent with the central role of UGT1A1 in bilirubin conjugation (e.g., Gilbert syndrome or subclinical phenotypes)^70^. In addition, UGT enzymes and CYP3A4 are inducible by medications and dietary supplements^71,72^. With limited information on such exposures, we cannot determine whether this population is the result of induction by medications or supplements.

In exploring the age-related changes in gene expression we were powered by both the number of donors and the expanded cell annotation of HLiCA. We were therefore able to explore age effects in the relatively rare mucus secreting cholangiocytes and identified an association between age and mucus production. Mucus production and mucus producing cholangiocytes have been described previously^2,49,73^ but it has not previously been shown that mucus production by intrahepatic cholangiocytes changes with age. In the intestine, mucus barriers are thinner and less functional with age^74,75^. While mucus barrier loss has not been described in the liver, here we observed increased expression of mucus production pathways, which may represent a compensatory response to age-related declines in hepatic mucus barrier integrity. Mucus in the liver protects cholangiocytes from damage caused by direct exposure to bile^73,76,77^, suggesting that maintenance of this barrier would be important throughout life. Although studies have focused on extrahepatic ducts, bile duct diameter has been reported to increase with age^78^. As we observed, mucus-secreting cells were enriched in larger intrahepatic ducts. The increased expression of mucus secretion genes we observed with age in the HLiCA may reflect age-related duct enlargement.

### HLiCA is built on the collective cell mapping and annotation efforts of many individuals and research groups, bringing these data together into a comprehensive atlas

Individual maps have often focused on specific cell lineages, whether enriching for them in the sample preparation or focusing data analysis efforts toward higher depth annotations of some lineages. The HLiCA annotation leverages information about expected cell types and markers from these individual maps to characterize the human liver cell types to the highest resolution currently possible. A crucial step in obtaining this cross study cell type resolution was the uniform read alignment (i.e to the same genome version with the same alignment software version) and the careful selection of an integration method. Another crucial resource was the iterative, community-based discussion of cell type annotations and corresponding marker genes. This harmonized data processing and collaborative cell annotation approach led to the identification of 47 cell types, expanding on the number of cell types identified in any individual map^2^. Previous efforts to integrate sc/snRNA-seq liver maps, used the existing cell type labels from the individual studies, leading to only a high level annotation of 20 cell types^79^. Our atlas presents the first ever atlas with refined cell annotations based on the integrated atlas.

### The HLiCA can be extended in the future as more samples, new data modalities and an even greater depth of donor clinical information become available

Although the half million cells included in the HLiCA is the largest liver atlas available to date it is possible we have still not identified all possible cell types in the healthy liver. For example, innate lymphoid cells (ILCs) are known to represent 1% of liver lymphocytes^80,81^. While liver ILCs have been observed using protein markers, there are no established gene expression markers for liver ILCs and we could not annotate a population in the HLiCA. For populations such as ILCs, combining gene expression with additional modalities, such as protein quantification from the same cells, could help define expression markers and clustering strategies for identifying these populations in the HLiCA or future atlases. The HLiCA was limited to 10x Genomics assays. Future atlas efforts could incorporate additional sc/snRNA-seq platforms as more data become available, enabling cross-assay integration. At the time of data collection, non-10x Genomics datasets were scarce, and including them would have introduced batch effects that would have been difficult to resolve. This analysis was also limited in the demographic data we could explore for gene expression associations. The liver cell atlas roadmap^17^ outlined a process for the harmonized collection of additional meta data to move beyond the age and sex analysis shown here. Additional information, such as time of sample collection, would be important as circadian rhythm is estimated to impact the expression of 22% of genes in the liver^82^. Standardization of sample information as outlined in the roadmap should enable the collection of relevant clinical information (including diet, basic metabolic index, alcohol and substance use) which could then be explored in future atlas versions. As more samples become available, the HLiCA could be further improved by incorporating tissue from healthy living donors^83,84^ as both tumour-adjacent liver tissue^85^ and tissue from deceased donors^86^ can exhibit altered gene expression profiles. Here we explored the overall diversity in genetic ancestry across the HLiCA and, by integrating data from multiple research centres, we improved on the representation of genetic ancestries compared to individual maps. However, there is a confounding effect between genetic ancestry representation and the research centre of origin, which in turn is confounded by technical effects such as cell enrichment and isolation protocols. It was therefore challenging in the HLiCA to associate gene expression with genetic ancestry independent of influence from technical effects. As there are known differences in liver drug metabolism between genetic ancestry populations^87^ and genetic variability in key marker genes used here^88^, it will be important for future liver cell atlases explore the dynamics of different cell types in relation to genetic ancestry

### Just as the HLiCA is built upon previously published maps of the human liver, our hope is that HLiCA will form a foundation for further atlassing of the human liver

We anticipate a major component of future atlases of the human liver going forward will focus on spatial transcriptomics. Additional cell types may exist in spatially distinct regions, which may have unique transcriptional profiles that cannot be teased apart with current data. We anticipate HLiCA v2.0 will be heavily influenced by the increasing availability of spatial transcriptomics data, to better understand how features including zonation, vasculature beds, and connective tissue influence cellular activity and crosstalk. With the release of this version of HLiCA we welcome feedback on the cell annotations, as knowledge of the cell types in the liver continually expands, as this will allow us to continue to improve the atlas. We anticipate the Cell Annotation Platform (CAP) will be a great place to house this feedback where it can be publicly discussed and built upon for HLiCA v2.0. The HLiCA v1.0 is designed to evolve with the field, integrating increasingly available spatial insights and community expertise to refine our understanding of human liver biology.

## Data Availability

All processed data generated in this study are available through the Human Cell Atlas (HCA) Data Portal at: https://data.humancellatlas.org/hca-bio-networks/liver and CELLxGENE at: https://cellxgene.cziscience.com/collections/059202e1-1f1b-483f-9151-f3a25a380c39. Raw sequencing data used in this study were generated by previously published studies and are available through their original repositories. Newly generated unpublished data from this study have been deposited in HCA Data Portal at: https://explore.data.humancellatlas.org/projects/cea413af-79b3-4f11-8b48-383fe9a65fbe. Code to reproduce all analyses, together with documentation for applying HLiCA as a reference atlas, is available at: github.com/quon-titative-biology/HLiCA.

All Supplementary Data are provided as a compressed archive and are additionally archived on Zenodo at: doi.org/10.5281/zenodo.20380927. Pretrained scANVI reference models^89,90^ (Supplementary Data 12-18) are available at: doi.org/10.5281/zenodo.20380927 to facilitate reuse of HLiCA as a reference atlas.

## Acknowledgements

We thank the research participants and their families for their invaluable contributions. The authors would like to thank Erica Marie Rutherford and Jason Hilton from the Lattice Team for their support with data management. Selected conceptual figures were created using BioRender.com. This publication is part of the Human Cell Atlas–www.humancellatlas.org/publications/.

## Author Contributions

**R.D.E.** contributed Data curation, Formal analysis and Biological interpretation; **Y.C. and H.H.** contributed Data curation and Formal analysis; **J.R.P.** contributed Data curation, Formal analysis and Biological interpretation; **D.P.** contributed Formal analysis and Methodology; **R.R.** contributed Data curation and Formal analysis; **D.S.** contributed Formal analysis and Biological interpretation; **D.R.N.** contributed Biological interpretation and Methodology; **T.T. and E.A.K.D.** contributed Formal analysis and Biological interpretation; **J.A., Z.A.C., R.G., S.K., Z.L., L.M., A.Mur., D.N., N.S., S.T., K.B.-H., E.I.L., R.L., N.R., O.Y., B.J.A., A.G.C., S.I., G.M.L., K.R.M., R.E.S., A.S., S.A.T., L.V. S.S.H and B.W.** contributed Biological interpretation; **R.N. and M.S.** contributed Formal analysis; **J.R.W.-K.** contributed Biological interpretation and Cohort recruitment and data generation; **T.A. and J.E.P.** contributed Methodology; **N.C.H.** contributed Conceptualization, Data curation, Biological interpretation and Cohort recruitment and data generation; **R.D., D.G., M.G., S.A.M., C.L.S., A.Mul., G.Q. and G.D.B.** contributed Conceptualization, Data curation and Biological interpretation. **All authors** contributed to Writing - review & editing.

