## supplementary for "HLiCA: An integrated cell atlas of the healthy human liver"

#### Table of Contents

Supplementary Methods and Tables

Supplementary Results

Supplementary Tables

Supplementary Data Descriptions (large tables provided separately)

### Supplementary Methods and Tables

#### Table S1. Summary of datasets included in the HLiCA.

| Research Centre | Samples | Donors | Assays | Suspension Type | Sample Type | Enrichment /depletion | Cells (after QC) | Study |
|---|---|---|---|---|---|---|---|---|
| Edin | 29 | 17 | 10x 3' v3 | cell | Healthy non-lesional liver tissue from patients undergoing undergoing surgical liver resection | Combinations: of CD45, EPCAM,CD144, ICAM2 | 92,193 | Unpublished |
| GIS/A*STAR | 23 | 23 | 10x 3' v3 | cell | Core needle biopsies from patients with chronic HBV infection or functionally cured HBV | Centrifugation–based parenchymal and non-parenchymal cell enrichment | 43,318 | Narmada et al J. Hep 2024 |
| JMU | 4 | 4 | 10x 3' v3 | cell | Healthy adjacent tissue from patients receiving a lobectomy or hemihepatectomy for oncologic resection | | 26,404 | Thomann et al. Nat. Com. 2026 (only 4/6 published samples available during HLiCA data collection) |
| Mass | 23 | 23 | 10x 3' v3 | nucleus | Healthy margins of excess surgical tissue | NA | 112,703 | Watson et al Nat. Com. 2025 |
| UHN | 40 | 24 | 10x 3' v2; 10x 3' v3; 10x 5' v1 | cell; nucleus | Caudate lobe from neurologically deceased donors | NA | 198,527 | MacParland et al Nat. Comm. 2018; Andrews et al Hep. Comm. 2022; Andrews et al J. Hep 2024 |
| VIB-UGent | 37 | 19 | 10x 3' v2; 10x 3' v3 | cell; nucleus | Biopsies from patients undergoing cholecystectomy or gastric bypass and healthy adjacent tissue removed during liver resection due to colorectal cancer metastasis | Combinations of: CD45+,CD45-,CD14+,CD16+ | 51,554 | Guilliams et al Cell. 2022 |
| | **156** | **110** | | | | | **524,699** | |

### Participant Information and Sample Processing from Research Centre: Edin

Local approval for procuring human liver tissue for scRNA-seq was obtained from the NRS BioResource and Tissue Governance Unit (study number SR574), following review at the East of Scotland Research Ethics Service (reference 15/ES/0094). All subjects provided written informed consent. Healthy non-lesional liver tissue was obtained intraoperatively from patients undergoing surgical liver resection for solitary colorectal metastasis, autosomal dominant polycystic liver disease or hepatocellular adenoma at the Hepatobiliary and Pancreatic Unit, Department of Clinical Surgery, Royal Infirmary of Edinburgh. Patients with abnormal liver function tests or those who had received systemic chemotherapy within the past 4 months were excluded from this cohort.

**Liver Digest**

Healthy human liver tissue was minced with scissors and digested in 5 mg ml−1 pronase (Sigma-Aldrich, P5147-5G), 2.93 mg ml−1 collagenase B (Roche, 11088815001) and 0.019 mg ml−1 DNase (Roche, 10104159001) at 37 °C for 30 min with agitation (200–250 r.p.m.), then strained through a 120-μm nybolt mesh along with PEB buffer (PBS, 0.1% BSA and 2mM EDTA) including DNase (0.019 mg ml−1). Thereafter, all processing was done at 4 °C. The cell suspension was centrifuged at 400g for 7 min, supernatant removed, cell pellet resuspended in PEB buffer and DNase added (0.019 mg ml−1), followed by additional centrifugation (400g, 7 min). Red blood cell lysis was performed (BioLegend, 420301), followed by centrifugation (400g, 7 min), resuspension in PEB buffer and straining through a 35-μm filter. Following another centrifugation at 400g for 7 min, cells were blocked in 5% human TruStain (BioLegend 422302) for 10 min at 4 °C before antibody staining.

**Table S2. Liver Digest Reagents**

| Source | Product | Catalogue Number |
|---|---|---|
| Roche | Collagenase B | 11088807001 |
| Sigma Aldrich | Pronase | P5147 |
| Roche | DNAse 1 | 10104159001 |
| Gibco | dPBS | 13492609 |
| MACS | BSA stock solution | 130-091-376 |
| Sigma Aldrich | Ethylenediaminetetraaceticdisodium salt dihydrate | E5134-500G |
| Biolegend | Red blood cell lysis buffer | 420301 |
| Biolegend | Human TruStain FcX | 422302 |

**FACS**

Incubation with primary antibodies was performed for 20 min at 4 °C. All antibodies, conjugates, lot numbers and dilutions are listed below in Table 1. After antibody staining, cells were washed with PEB buffer. Cell viability staining (DAPI; 1:1,000) was performed, immediately before sample acquisition of Endothelial and Mesenchymal cell populations.

Human cell sorting for scRNA-seq was performed on a BD FACSAria™ Fusion. Viable single EPCAM negative, CD102/CD144 positive (Endothelia) or CD102/CD144 negative, CD45 negative (Mesenchyme) cells were sorted from human liver tissue.

**Table S3. Flow Antibodies**

| Name | Source | Catalogue Number | Lot Number | Dilution |
|---|---|---|---|---|
| EPCAM | Biolegend | 324232 | B292867 | 1 in 100 |
| CD45 | Biolegend | 304016 | B340163 | 1 in 100 |
| CD144 | BD Biosciences | 561566 | 2052834 | 1 in 100 |
| CD102 | Biolegend | 328506 | B293610 | 1 in 100 |
| HLA-DR | Biolegend | 307650 | B235468 | 1 in 100 |

**Gating Strategy**

Cells: FSC-A/SSC-A
F Singlets: FSC-H/FSC-A
S Singlets: SSC-H/SSC-A
Live cells: SSCA-A/DAPI-ve
Endothelial cells: EPCAM-ve, CD102/CD144+ve
Leucocytes: CD102/CD144-ve, CD45+ve/HLA-DR+ve cells
Mesenchyme: CD102/CD144-ve, CD45-ve cells

**scRNA-seq Processing**

Single cells were processed through the Chromium Single Cell Platform using Chromium Single Cell 3′ Library and Gel Bead Kit v3 and v3.1 (10X Genomics, PN-1000075 and PN-1000121), and the Chromium Single Cell B Chip Kit (10X Genomics, PN-1000073) and Chromium Single Cell G Chip Kit (10X Genomics, PN-1000120) as per the manufacturer's protocol. In brief, single cells were sorted into PBS plus 0.1% BSA, washed twice and counted using a Bio-Rad TC20. Then, 10,800 cells were added to each lane of the 10X chip. The cells were partitioned into Gel Beads in Emulsion in the Chromium instrument, in which cell lysis and bar-coded reverse transcription of RNA occurred, followed by amplification, fragmentation and 5′ adaptor and sample index attachment. Libraries were sequenced on an Illumina HiSeq 4000 or NovaSeq6000.

#### Sample Sources and Collection

Samples were obtained from a range of tissue sources, including biopsies, surgically resected liver tissue, and whole liver lobes (**Table S1**). Biopsies and resected tissues were collected from living donors, whereas whole lobes were obtained from neurologically deceased donors without known liver disease. The sample type and collection method mainly varied between Research Centre but were consistent within (**Table S1**). We therefore evaluated whether data from multiple Research Centres were contributing to the described cell types as well as age and sex signals, to control for potential technical signals from sample type.

Clinical information was limited and not uniformly collected across research centres. A summary of the available data is provided in (**Extended Data Table S5**). However, considerable differences in data collection were observed between Research Centres.

#### Sex confirmation from gene expression

Sample sex was assessed using expression of the sex chromosome–linked genes *XIST* and *RPS4Y1*. For each sample, we calculated the median and 75th percentile (Q3) expression of these genes across all cells. Samples with Q3 > 0 for *XIST* and not *RPS4Y1* were classified as female, while those with Q3 > 0 for *RPS4Y1* and not *XIST* were classified as male. Three samples with discordant or ambiguous expression patterns were excluded from sex and age analyses due to concerns regarding potential metadata misannotation.

#### Integration with pediatric liver

To evaluate the utility of the HLiCA as a reference for human liver cell types we integrated with a smaller (n=7) pediatric liver map. We explored the lymphocyte and mesenchyme lineages in both data sets for shared cell types. We integrated HLiCA and pediatric data for both lineages with Harmony (v1.2.3) with the following parameters: group.by.vars set to the library_alias (sample ID in pediatric data), assay.use set to 'RNA', and reduction set to 'pca'. Clusters in both the lineages were defined using the Seurat functions "*FindNeighbours*" and "*FindClusters*" on the first 30 harmony dimensions.

### Supplementary Results

#### Integration benchmarking results

All four integration methods performed comparably across metrics (**Fig. S1**). Most approaches improved the mixing of cell types relative to no integration, as indicated by lower cell type ARI and higher cell type LISI, with the expectation of WNN. All methods improved the mixing of suspension type and library, with stronger library mixing generally associated with greater cell type mixing.

Harmony produced the most conservative integration, yielding the lowest ARI values across categories. As the expert review of members from the bionetwork did not indicate over-integration, Harmony was selected for downstream analyses due to its strong performance in integrating libraries and suspension types.

#### Integration with pediatric liver

In a map of human pediatric liver map (n=9 donors; n= 42,660 cells) several cell types identified in HLiCA were not seen, including *NRXN1+* stromal cells and regulatory T cells. When looking at the pediatric mesenchyme, the lineage where *NRXN1+* stromal cells were found in HLiCA, we do see a cluster of 85 cells with high *NRXN1* expression (**Fig. S11**). However, 81% of these cells are from one pediatric donor (**Fig. S11**), making it challenging to confidently assign this population as a unique cell type and individual donor specific population. When integrated with the HLiCA, these cells cluster with *NRXN1+* stromal cells, suggesting they are real and lending confidence that these cells can be found in the pediatric liver as well.

In the case of T regs which were annotated in HLiCA but not the pediatric map, we do not see any unique cluster of potential T regs in the pediatric map alone based on the key marker *IL2RA* (**Fig. S9**). When clustered with the HLiCA there is a population of 69 pediatric cells clustering with the HLiCA T regs, suggesting there were potentially T regs in the pediatric samples but there were not enough of them or with a unique enough cell type signature to clusters separately in the pediatric samples alone (**Fig. S9**).

#### Pretrained scANVI reference models

We trained scVI models using scvi-tools on the entire HLiCA atlas as well as each lineage. scVI was applied to each dataset with batch effects accounted for similar to as in the Harmony integration. Specifically, with the following parameters: batch_key set to 'donor_uuid' and with categorical_covariate_keys set to 'STUDY', 'suspension_type', 'assay', and 'library_alias'. To enable the use of the HLiCA as a cell annotation reference, scANVI was trained on the cell type labels. All models were trained using default parameters. Resulting models are provided in **Supplementary Data 12-18**.

### Supplementary Data Descriptions

*Large tables provided as csv files at: doi.org/10.5281/zenodo.20380927*

**Supplementary Data 1**: The file "HLiCA_sample_metadata.csv" contains sample **metadata** used in the HLiCA analysis.

**Supplementary Data 2**: The file "HLiCA_extended_metadata.csv" contains **extended metadata** compiled from original publications where available.

**Supplementary Data 3:** The file "HLiCA_lineage_markers.csv" contains **lineage-level marker genes** used for annotation of major cell populations in the HLiCA atlas. Markers are categorized as canonical, based on established literature, or additional, based on feedback from the Bionetwork during HLiCA construction.

**Supplementary Data 4**: The file "HLiCA_cell_type_markers.csv" contains **cell type-specific marker genes**. Genes are categorized by direction of expression as positively associated (POS) or negatively associated (NEG) with each cell type. These markers are intended for distinguishing cell types within a given lineage.

**Supplementary Data 5:** The file "HLiCA_CellType_DEG.csv" contains cell type-specific **differential expression results**. Only differentially expressed genes (FDR < 0.005 and $|\log_2$ fold change$| > 1$) are included. Statistics compared a gene expression in the listed cell type to all other cell types in the lineage.

**Supplementary Data 6:** The file "HLiCA_CellType_GSEA.csv" contains cell type-specific **GSEA** based on differentially expressed genes for the cell type. Significantly enriched pathways (Padj < 0.05) are shown. Reported statistics include nominal p-value, Benjamini–Hochberg adjusted p-value (Padj), enrichment score (ES), normalized enrichment score (NES), gene set size after filtering, and leading-edge genes contributing to the enrichment signal.

**Supplementary Data 7**: The file "HLiCA_community_standard_cellnames.csv" contains Community-standard cell type names used to align the HLiCA annotations with **Cell Ontology** terms.

**Supplementary Data 8:** The file "HLiCA_Age_DEG.csv" contains cell type-specific **differential expression results** with **age**. Only differentially expressed genes (FDR < 0.05, |coef| > 0.5) are included. A positive coef indicates genes which increased in expression with age and a negative coef indicates genes which decreased in expression with age.

**Supplementary Data 9:** The file "HLiCA_Age_GSEA.csv" contains cell type-specific **GSEA** based on differentially expressed genes with **age** in the cell type. Significantly enriched pathways (Padj < 0.05) are shown. Reported statistics include nominal p-value, Benjamini–Hochberg adjusted p-value (Padj), enrichment score (ES), normalized enrichment score (NES), gene set size after filtering, and leading-edge genes contributing to the enrichment signal.

**Supplementary Data 10:** The file "HLiCA_Sex_DEG.csv" contains cell type-specific **differential expression results** with **sex**. Only differentially expressed genes (FDR < 0.05, |coef| > 0.5) are included. A positive coef indicates genes more highly expressed in females and a negative coef indicates genes more highly expressed in males.

**Supplementary Data 11:** the file "HLiCA_Sex_GSEA.csv" contains cell type-specific **GSEA** based on differentially expressed genes with **sex** in the cell type. Significantly enriched pathways (Padj < 0.05) are shown. Reported statistics include nominal p-value, Benjamini–Hochberg adjusted p-value (Padj), enrichment score (ES), normalized enrichment score (NES), gene set size after filtering, and leading-edge genes contributing to the enrichment signal.

**Supplementary Data 12-18:** The compressed folder "scANVI_models.zip" contains trained **scANVI models** for each HLiCA cell lineage as well as the entire atlas.

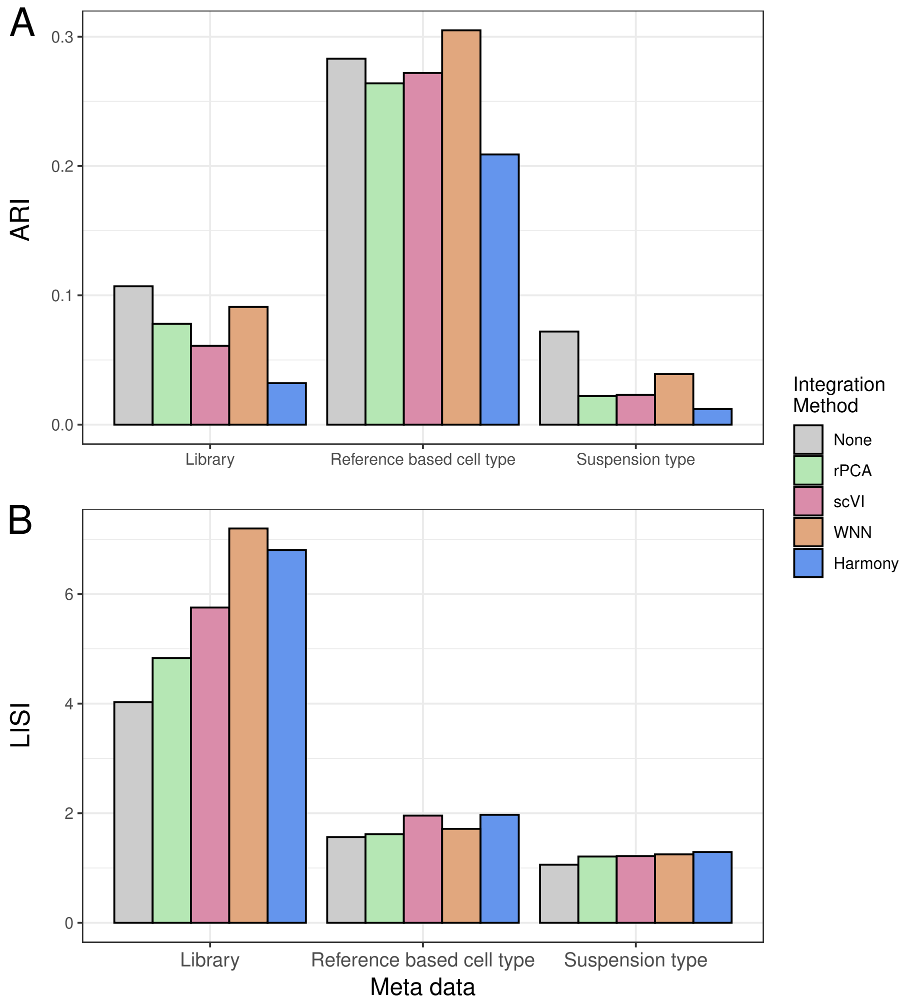


**Figure S1: Integration methods perform similarly for two metrics on mixing of three metadata variables.** A) Adjusted Rand Index (ARI) calculated for a reference-based cell type (not the annotations derived from the final integrated object), library (i.e., sample), and suspension type (i.e., single-cell or single-nucleus) following integration using four methods. B) Local Inverse Simpson Index (LISI) calculated for the mixing of the same metadata and methods as in A.

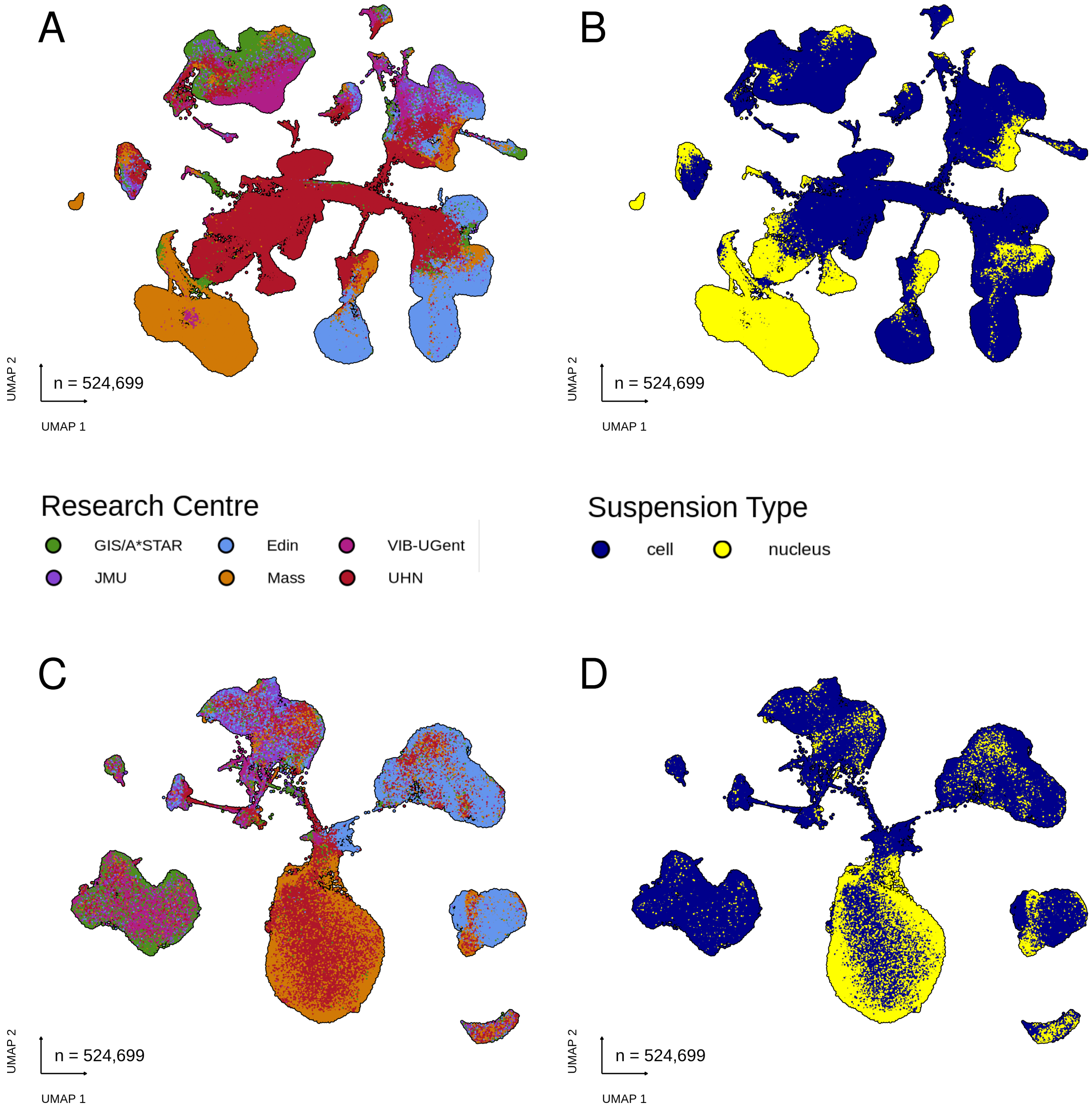


**Figure S2: UMAP of cells that passed quality control before and after Harmony integration.** A) UMAP of merged cells coloured by research centre. B) UMAP of merged cells coloured by suspension type. C) UMAP of integrated cells coloured by research centre. D) UMAP of integrated cells coloured by suspension type.

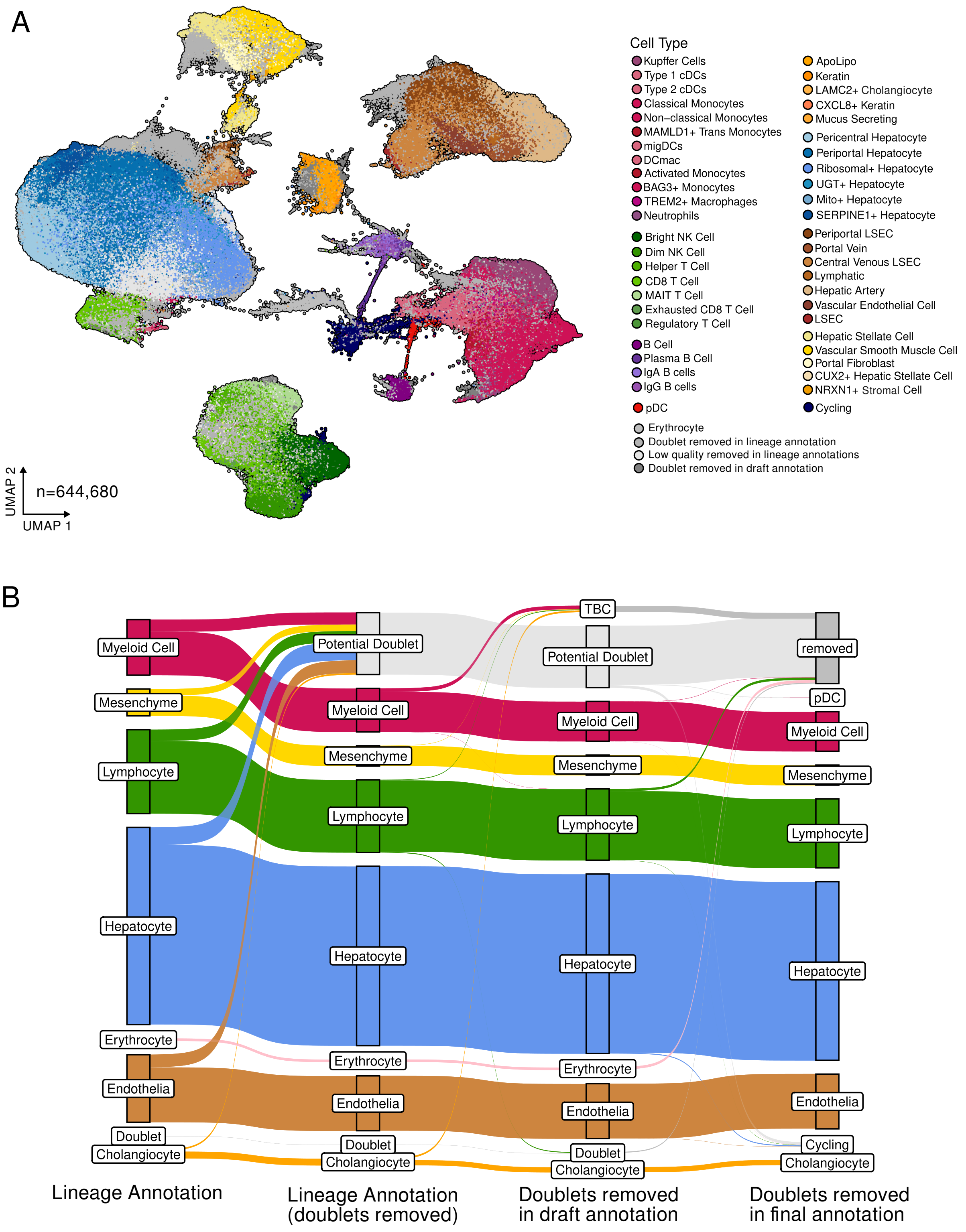


**Figure S3: Quality control process used to remove cells.** A) UMAP of the integrated HLiCA coloured by the final cell annotation, with cells considered doublets and erythrocytes included and coloured in grey. B) Sankey showing major cell type lineages and the removal of doublets from each lineage across stages of cell annotation.

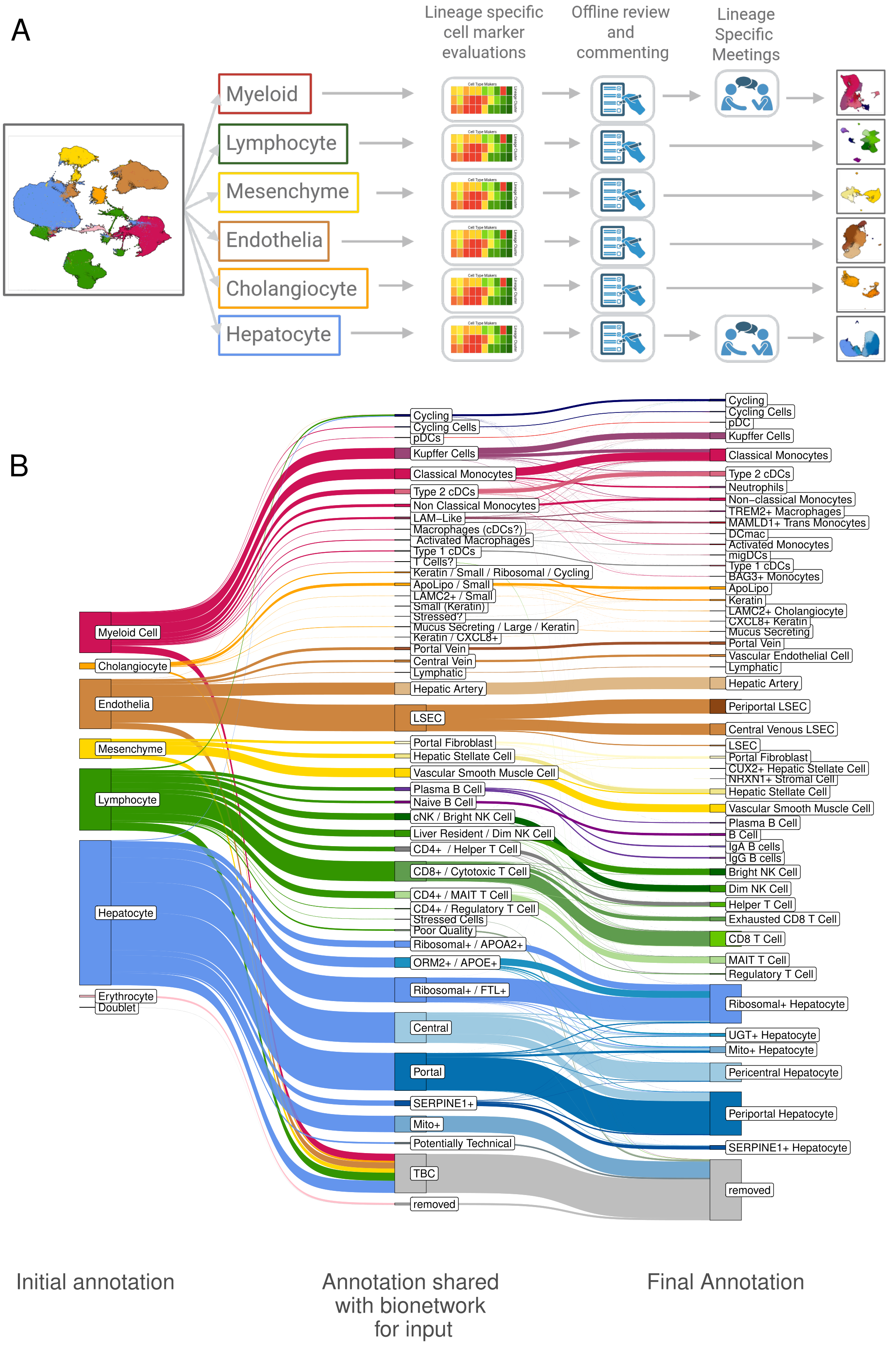


**Figure S4: Cell annotation process toward final Gamma annotation.** A) Cells were annotated within major lineages, with feedback provided offline and in lineage-specific meetings for further discussion. B) Sankey showing steps of cell annotation.

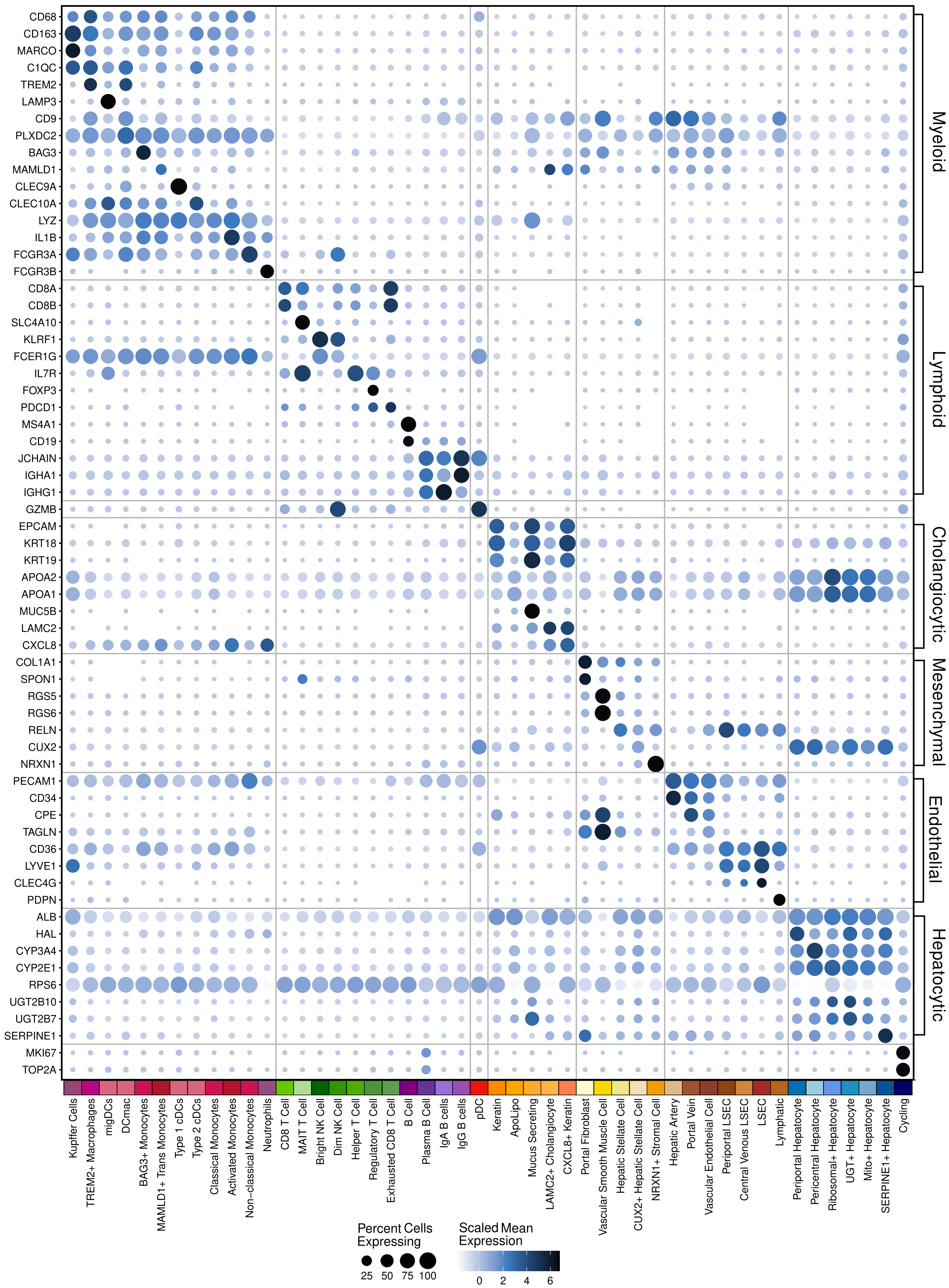


**Figure S5: Representative cell type markers used in the annotation process.**

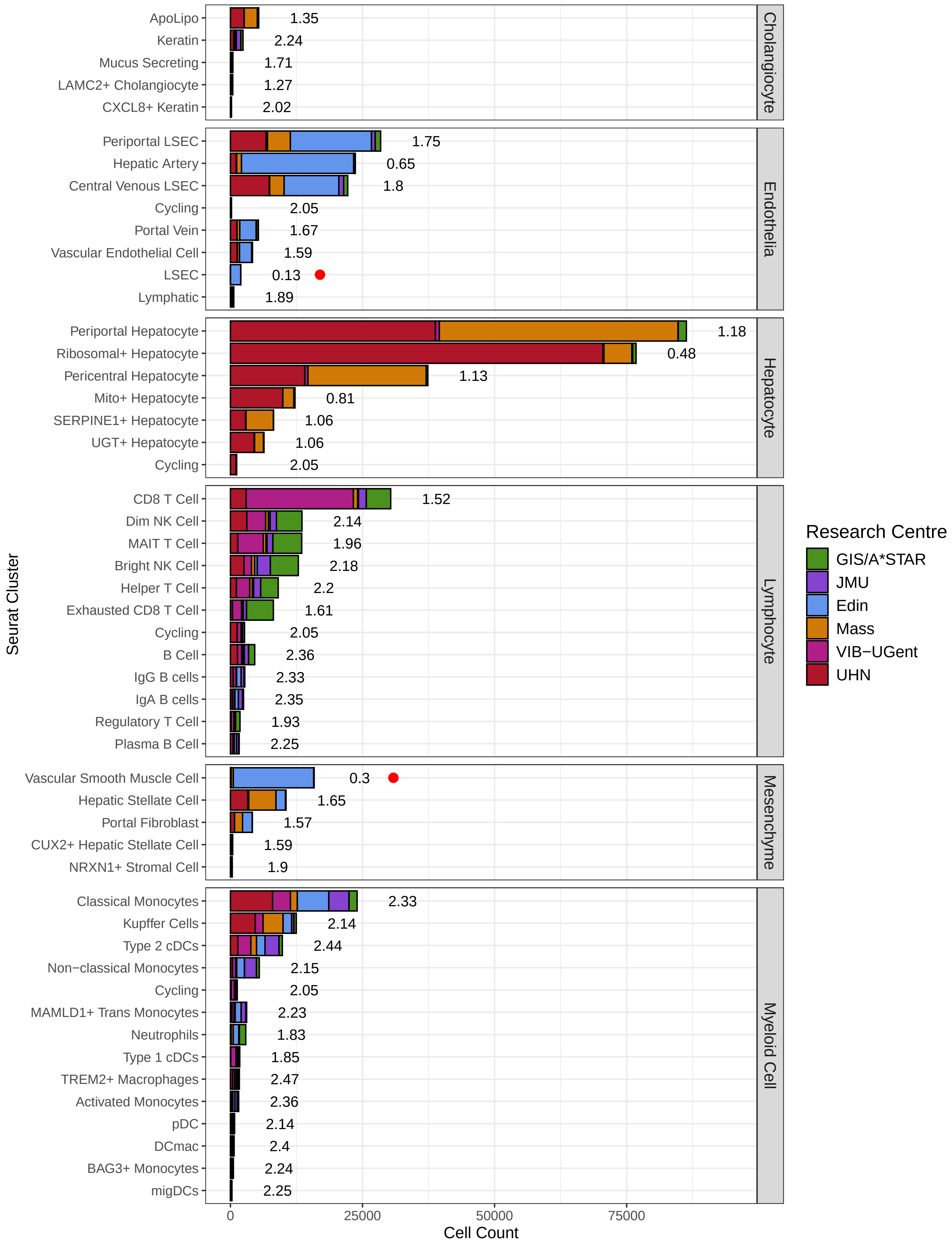


**Figure S6: Most cell types are well mixed by Research Centre.** A) Bars show cell counts from each Research Centre split by cell type. Numbers denote Shannon entropy of Research Centre mixing per cell type; higher values indicate greater mixing (maximum = 2.32). Red dots denote cell types failing the mixing threshold ($\geq$95% of cells from a single Research Centre).

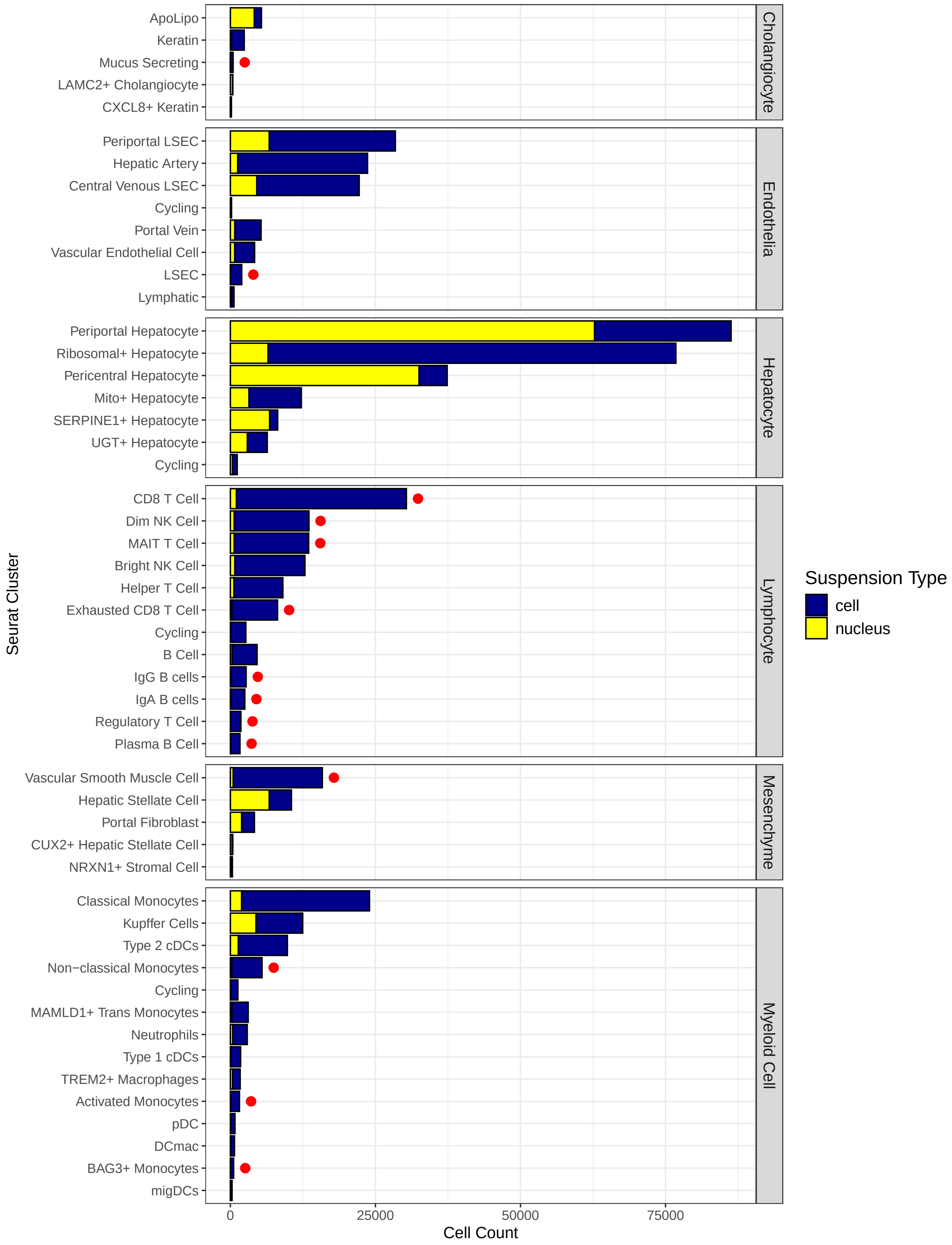


**Figure S7: Cell types are well represented in both suspension types (single-cell and single-nucleus), with enrichments following expected patterns.** A) Bars show cell counts from each suspension type split by cell type. Red dots denote cell types failing the mixing threshold (≥95% of cells from one suspension type). The entire atlas consists of 70% single-cell and 30% single-nucleus data.

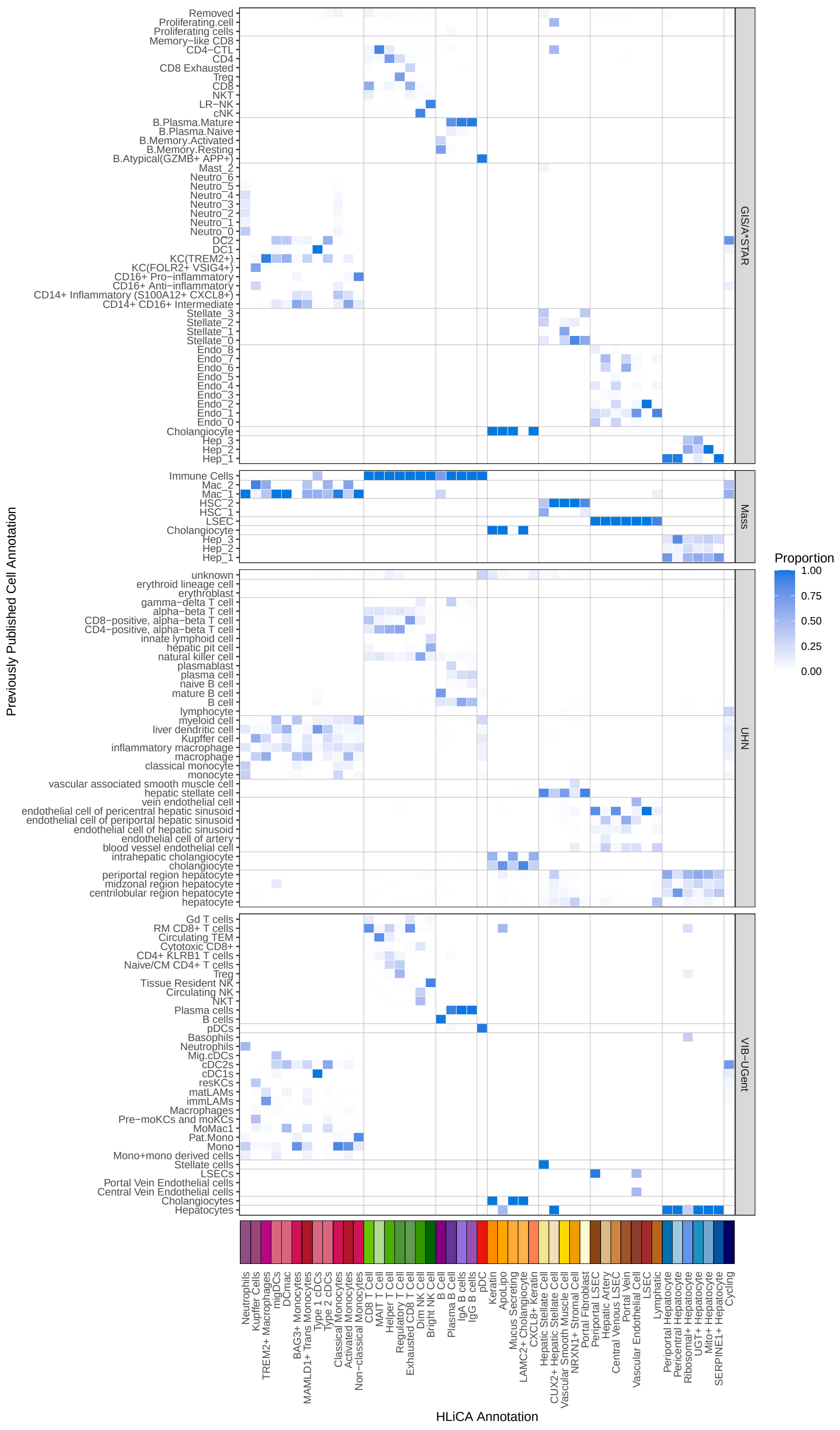


**Figure S8: Confusion matrix comparing final HLiCA annotations with previously published cell labels for annotated cells.** Proportions indicate, for each Gamma annotation, the fraction of cells originating from each original label.

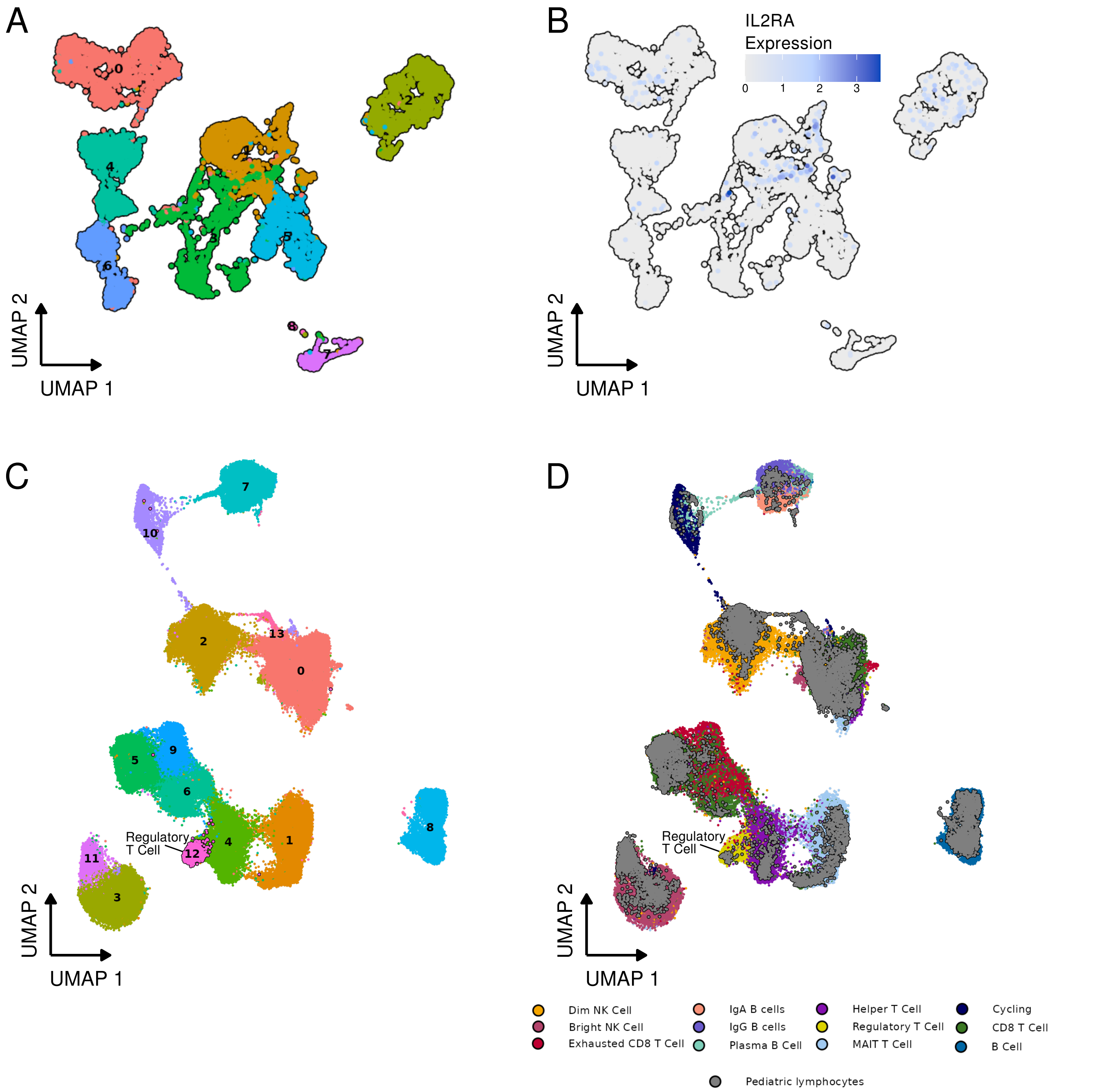


**Figure S9: Regulatory T cells can be seen in the pediatric liver when integrated with the HLiCA.** A) Pediatric lymphocyte UMAP with points coloured by cell type cluster. B) IL2RA (regulatory T cell marker) expression in pediatric lymphocytes. D) Integrated UMAP of pediatric and HLiCA lymphocytes coloured by cluster, with regulatory T cells outlined in black. E) Integrated UMAP of pediatric and HLiCA lymphocytes coloured by HLiCA cell type annotation, with pediatric cells shown in grey and outlined in black.

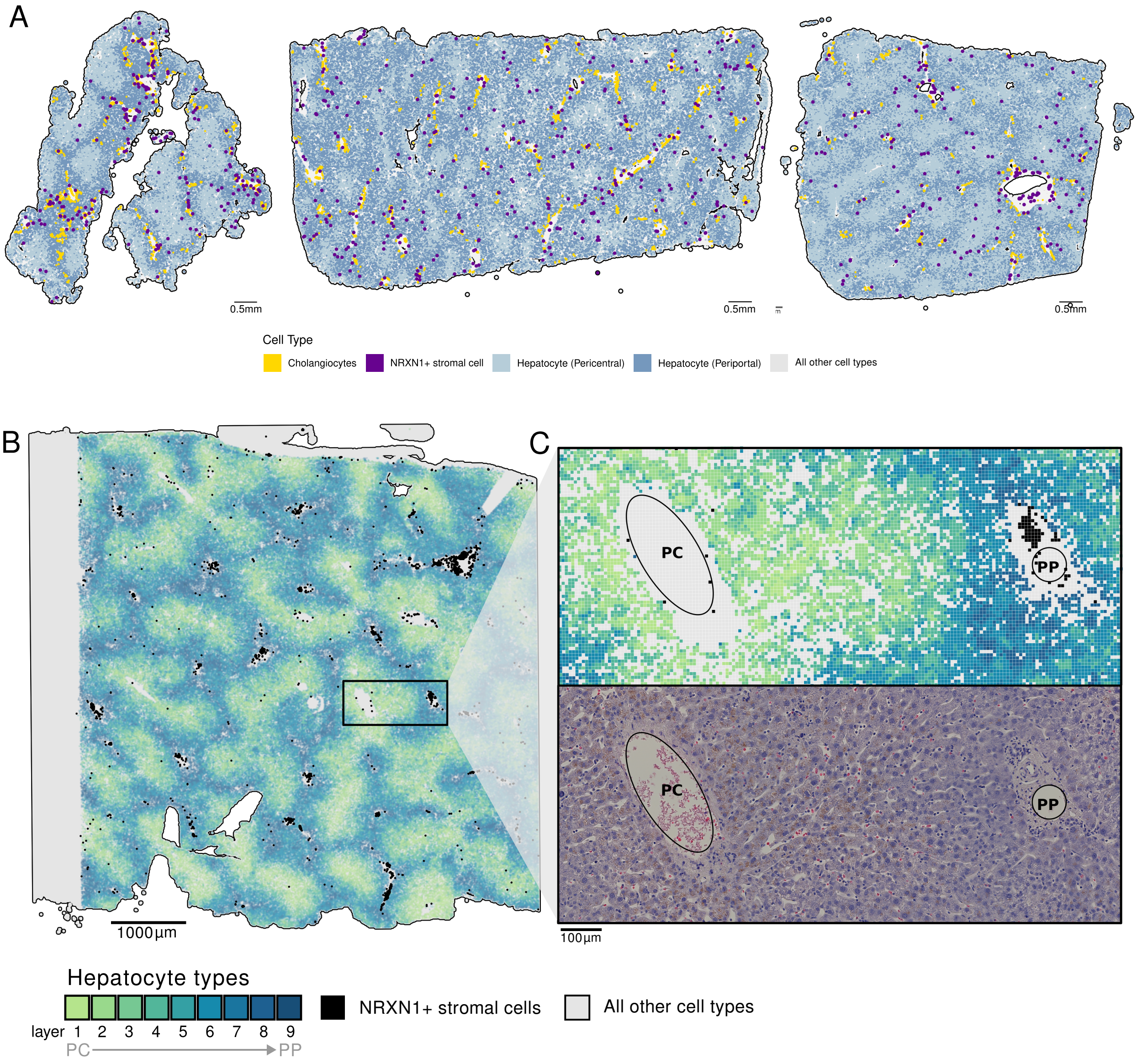


**Figure S10: NRXN1+ stromal cells were observed in Xenium and VisiumHD spatial transcriptomics and are more abundant in periportal regions.** A) Location of NRXN1+ stromal cells in Xenium spatial transcriptomics. Only selected cell type annotations are shown to highlight zonation patterns. B) Zonation of hepatocytes in VisiumHD data, with only hepatocytes shown to highlight zonation patterns. NRXN1+ stromal cells are shown in black. C) Zoomed view of a PC-to-PP region with 8 μm bin annotations of hepatocytes and NRXN1+ stromal cells shown. H&E images are aligned and displayed to assess liver structural features.

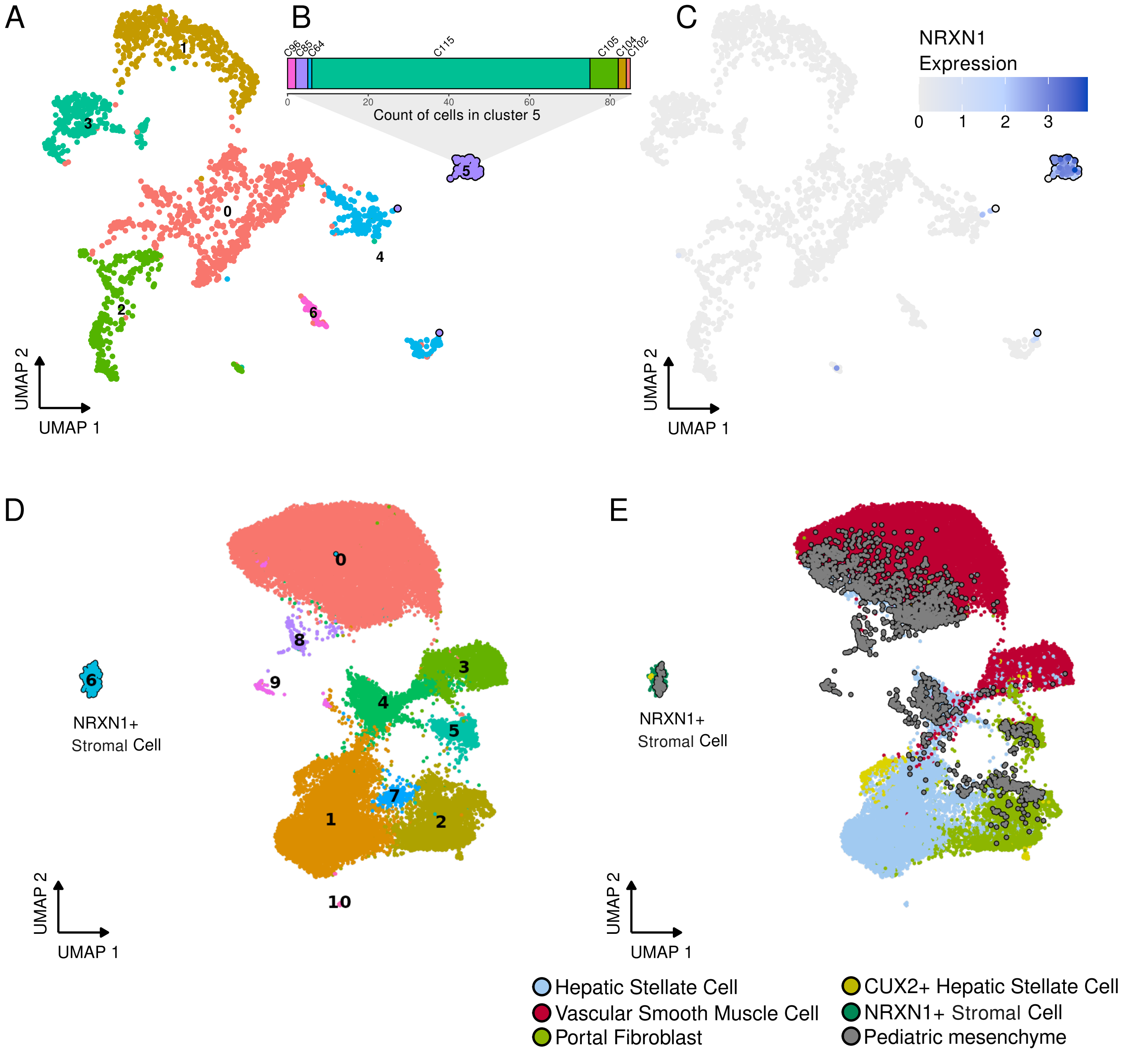


**Figure S11: NRXN1+ stromal cells can be seen in the pediatric liver when integrated with the HLiCA.** A) Pediatric mesenchyme UMAP with points coloured by cell type cluster. B) Contribution of seven donors to the NRXN-high cluster. C) NRXN1 expression in pediatric mesenchyme. D) Integrated UMAP of pediatric and HLiCA mesenchyme coloured by cluster, with NRXN1+ stromal cells outlined in black. E) Integrated UMAP of pediatric and HLiCA mesenchyme coloured by HLiCA cell type annotation, with pediatric cells shown in grey and outlined in black.

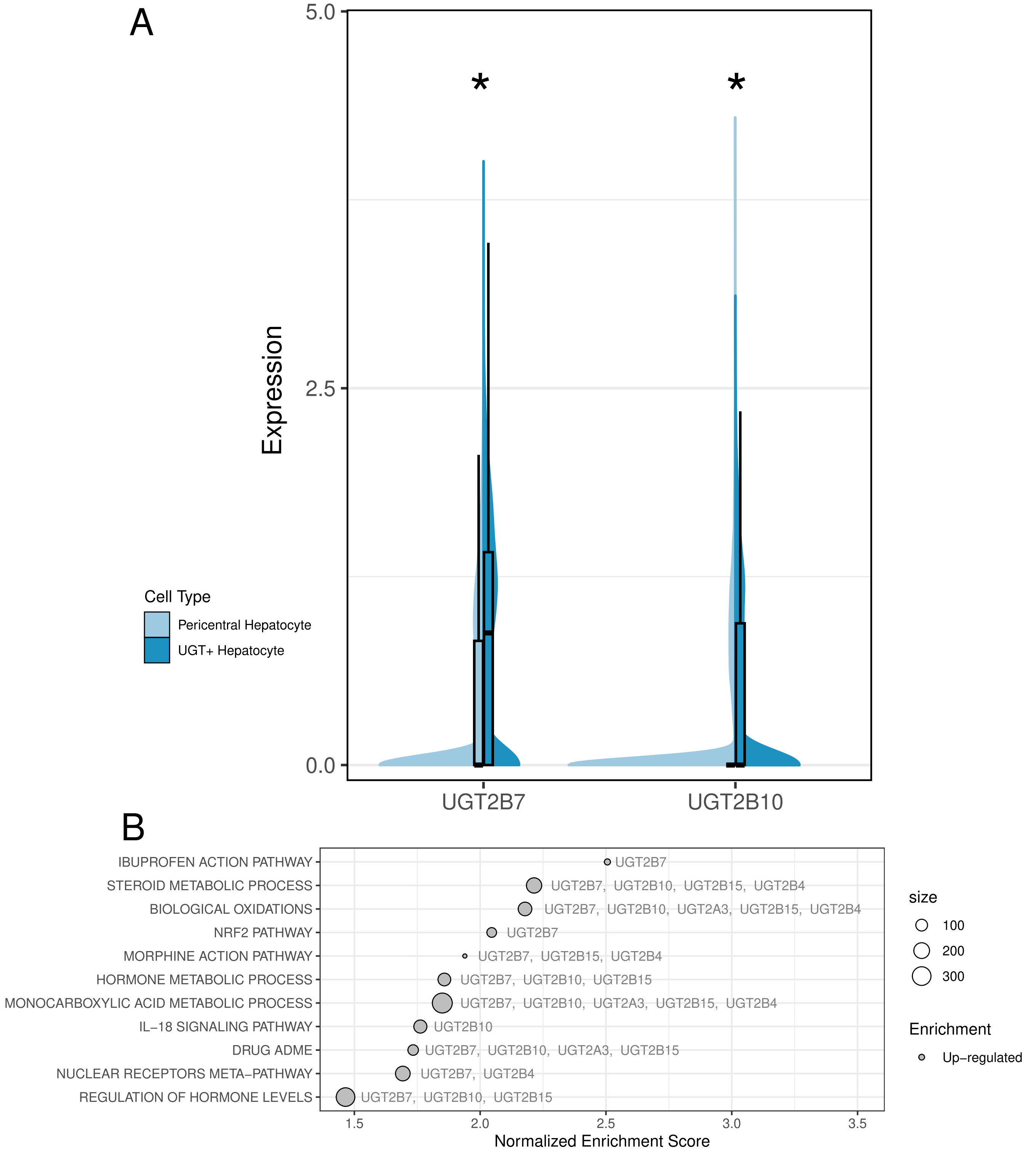


**Figure S12: UGT+ hepatocytes exhibit higher UGT gene expression and pathway enrichment compared to pericentral hepatocytes.** A) Expression of two UGT genes differentially expressed between UGT+ and pericentral hepatocytes. B) GSEA of UGT-related pathways. All shown pathways include UGT genes and are enriched in UGT+ hepatocytes relative to pericentral hepatocytes.

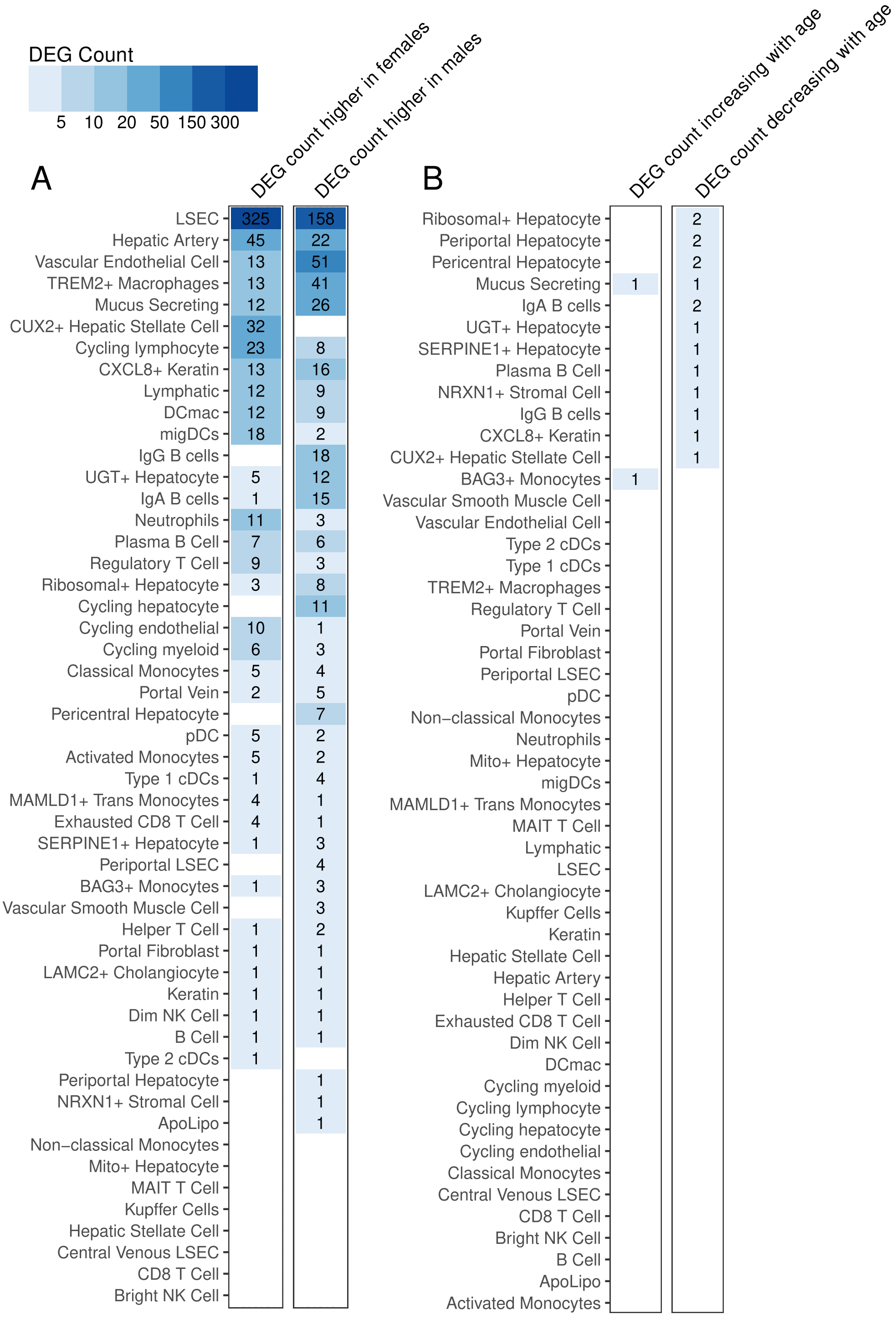


**Figure S13: Counts of differentially expressed genes (DEGs) in each cell population.** A) Sex-associated DEGs. B) Age-associated DEGs.

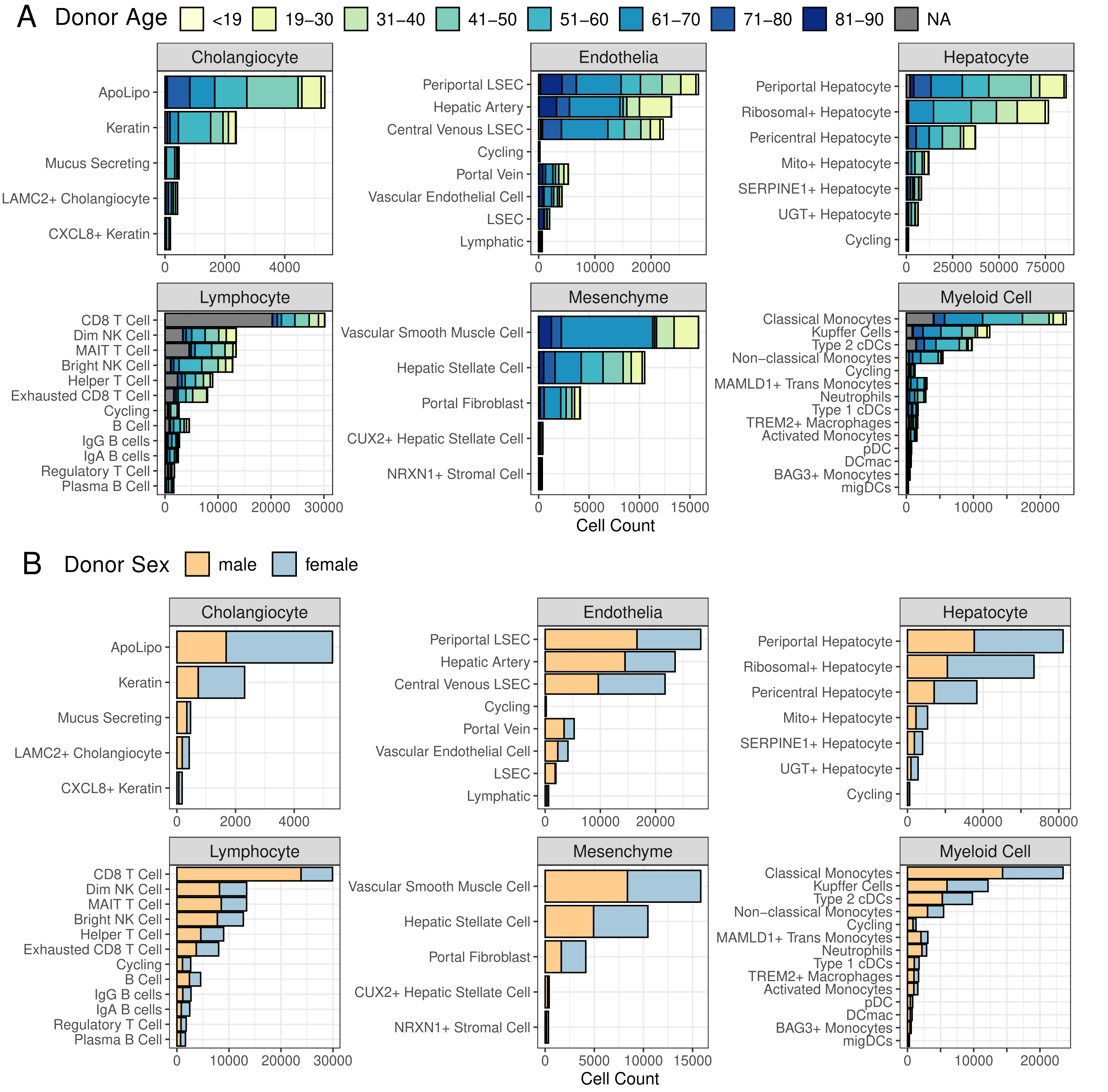


**Figure S14: Counts of each cell type with bars coloured by donor characteristics.** A) Age. B) Sex.
